# An Explainable Machine Learning Approach to study the positional significance of histone post-translational modifications in gene regulation

**DOI:** 10.64898/2026.01.30.702742

**Authors:** Sharun Ramachandran, Nithya Ramakrishnan

**Affiliations:** Institute of Bioinformatics and Applied Biotechnology, India

**Keywords:** Epigenetics, Histone PTMs, ChIP-seq, Machine Learning, SHAP

## Abstract

Epigenetic mechanisms regulate gene-expression by altering the structure of the chromatin without modifying the underlying DNA sequence. Histone post-translational modifications (PTMs) are critical epigenetic signals that influence transcriptional activity,
promoting or repressing gene-expression.Understanding the impact of individual PTMs and the combinatorial effects is essential to deciphering gene regulatory mechanisms.In this study,we analyzed the ChIP-seq data for 26 PTMs in yeast, examining the PTM
intensities gene-wise from positions-3 to 8 in each gene.Using XGBoost classifiers, we predicted gene transcription rates and identified
key histone modifications and nucleosomal positions that are critical in gene-expression using explainability measures (such as SHAP).
Our study provides a comprehensive insight into the histone modifications, their positions and their combinations that are most critical
in gene regulation in yeast.The proposed explainable Machine Learning models can be easily extended to other model organisms to
provide meaningful insights into gene regulation by epigenetic mechanisms.

## 1 INTRODUCTION

Gene-expression is regulated by epigenetic mechanisms that govern chromatin accessibility and transcriptional activity [1, 2].. Among known epigenetic factors, histone post-translational modifications (PTMs) are critical regulatory marks, determining whether a gene is actively transcribed or repressed [3, 4]. In yeast, similar to higher model organisms, PTMs are crucial for maintaining cellular identity and function [5, 6]. Histone proteins form the structural core around which DNA is wrapped, and the chemical modifications along the histones’ tails - such as acetylation, methylation, and phosphorylation — play pivotal roles in regulating gene-expression [1].

The positions of histone modifications across promoter regions, transcription start sites (TSS) and gene bodies provide important spatial context for their regulatory functions [7, 8]. For example, histone marks such as H3K4me3 and H3K9ac are highly active near promoters and TSSs of actively transcribed genes, showing their positional importance in transcriptional activation [9]. Similarly, it has been established that the combinatorial and positional distribution of PTMs across promoters and gene bodies correlates strongly with gene-expression levels [10]. Thus, it is not only the presence of the modification, but also its location within the chromatin landscape that regulates its influence on transcriptional effects. This spatial and combinatorial patterning of PTMs constitutes the histone code hypothesis, which suggests that specific combinations and arrangements of histone modifications transcriptional activity [4]. Studying the positional distribution and impact of histone PTMs across the genes is essential to unravel the complexities of gene regulation.

Recent studies in bioinformatics and machine learning (ML) have enabled quantitative modeling of the relationships between histone modifications and gene-expression. For example, ML models were used to predict gene-expression from histone modification profiles [11]. Additionally, multiple PTMs were integrated to improve prediction accuracy [12], while deep learning was applied to capture the dependencies between histone marks and transcriptional output [13]. These models not only capture linear associations, but also uncover non-linear interactions between modifications and their positional context, highlighting the importance of cross-play between marks. It was further demonstrated that these modifications affect nucleosome stability and higher-order chromatin organization [14], highlighting the combinatorial and positional inter-dependence among PTMs. Although the biochemical functions of individual modifications are relatively well-characterized, the rules governing their combinatorial and positional interplay are not adequately analyzed.

Several machine learning (ML)-based studies have been explored to examine the role of epigenetic modifications in regulating transcriptional activity and chromatin state dynamics [15, 16]. The approaches have demonstrated that PTMs carry information that can be computationally leveraged to understand gene-regulation mechanisms. Regression models were employed to predict PTM levels based on diverse genomic and biochemical features such as replication timing, nucleosome turnover rate, sense and antisense transcription levels, and histone-variant occupancy [17]. Similarly, support-vector regression was utilised to demonstrate that a combination of PTMs could accurately predict gene-expression levels, thus highlighting the predictive potential of histone marks in transcriptional regulation [18]. Other studies have integrated PTM data with DNA methylation and chromatin-accessibility features to improve transcriptional-outcome predictions [11].

Despite these advances, the identification of critical nucleosomal positions in the specific PTMs that influence gene regulation remains inconclusive [19–21]. While previous computational works have largely focused on modeling global relationships between epigenetic signals and gene-expression [11, 12, 18, 22], they have overlooked the positional dependencies and combinatorial interactions among histone PTMs across the chromatin. In this study, we overcome this gap by training Machine Learning models to predict gene-expression from histone modification patterns across nucleosomal positions, and subsequently employing SHAP (SHapley Additive exPlanations) values to interpret model predictions. Recent work by Chhatbar *et al* [23] employed deep learning models combined with SHAP to predict RNA Pol-II occupancy from chromatin-associated protein profiles under unperturbed conditions, demonstrating that SHAP importance scores can recover direct regulatory targets and capture the magnitude of transcriptional changes. Furthermore, SHAP explainer has also been applied to deep learning models for identifying key epigenetic predictors such as PTMs for gene-expression in humans [24]. However, they have not analyzed the critical positions/patterns of the PTMs in gene regulation. Additionally, SHAP has also been used to identify the crucial amino acid motifs in binding affinity prediction [25]. In our work, by quantifying the contribution of each PTM at each nucleosomal position, SHAP analysis helps us in identifying the most influential histone PTMs and their corresponding critical positions associated with transcriptional activity in yeast. This technique can be applied to different model organisms to compare the key PTM distributions across the nucleosomes regulating gene-expression [26].

## 2 MODELS AND METHODS

### 2.1 Data Preprocessing and Standardization

The nucleosome-level dynamics for 26 histone marks in yeast were captured at a high-resolution, genome-wide experiment recently [17]. In our work, we consider the steady-state values of the histone PTMs across the yeast genome, normalized to the H3 levels, at steady-state conditions, to predict the transcription rate of genes (obtained from [27]). The dataset for 26 PTMs [17] across all the genes were preprocessed so that each gene was represented as a 11-dimensional vector, with the *−*3 position to *−*1 representing the promoter nucleosomes and +1 to +8 representing the gene-body nucleosomes (considering the median length of an yeast gene to be 1.4 kbp [28]) - we therefore constructed a three-dimensional matrix of size 26 PTMs, *n* genes, 11 positions for the ML models. Standardization was performed using z-score normalization, transforming each feature (PTM) to have a mean of zero and a standard deviation of one.

### 2.2 Label Mapping and Thresholding

We employed the dataset from a study on transcriptional analysis on yeast for obtaining the labels for yeast gene [27]. In their work, high-density DNA microarrays were employed to quantify the abundance of mRNA for each yeast gene, using systematic perturbations of transcription initiation components. Normalized expression data were utilized to estimate relative transcription rates across multiple conditions, with mRNA abundance serving as a control for transcription rate under steady-state assumptions. This approach provided one of the first genome-wide maps of transcriptional regulation and gene-expression dynamics in eukaryotes.

To classify the yeast genes based on their transcriptional activity,the continuous mRNA measurements obtained from this study were transformed into categorical labels using a median expression threshold of 13 mRNA/hr. This median threshold was chosen considering similar previous studies on biological systems [29]) . Yeast genes with expression values above/below 13 mRNA/hr were categorized as high/low expressed (class 1/0).

However, one of the major issues with this approach was class imbalance between the low and high transcribing genes. Of the 4, 585 genes analyzed, approximately 3, 500 were categorized as high-expressed, leaving only 1,358 in the low-expression class. This imbalance led to biased and ineffective predictions, as models often favored the majority class. The challenges associated with class imbalance in genomic prediction tasks were emphasized in prior investigations [30], where it was demonstrated that skewed class distributions bias model performance towards dominant categories.

### 2.3 Oversampling using SMOTE

To address the imbalance between high-expression and low-expression gene classes, the KMeans-SMOTE (Synthetic Minority Oversampling Technique) algorithm was applied to the training data. SMOTE was first introduced as a method for synthesizing new minority-class samples by interpolating between existing ones, thereby improving classifier performance on imbalanced datasets [31] . In our work, KMeans-SMOTE was employed to improve the quality of synthetic samples by clustering the minority-class instances before generating new ones. This approach preserves the natural structure of the minority class and reduces the introduction of noisy or unrealistic samples. SMOTE was implemented using the `imblearn.over_sampling.SMOTE` module from the `imbalanced-learn` Python package (version ≥ 0.10). After applying KMeans-SMOTE, the ratio of genes in the two classes was balanced from 750:2700 to 2700:2700, ensuring an equal representation of both expression categories.

### 2.4 Machine Learning for gene-expression Prediction

To predict gene-expression levels from histone modification patterns, several machine learning (ML) models such as Logistic regression, XGBoost, SVM (Support Vector Machines) and Random forest classifier were evaluated. XGBoost demonstrated the highest predictive performance (See Supplementary Information (SI) Fig. S1). After grid-search was performed for selecting the best hyper-parameters (Sec. 2.5), a 10-fold cross-validation was performed, following which the means and standard deviations of the accuracies and F1-scores of the folds were assessed. The standard deviations were around 2% (Table. S1 of the SI) indicating that the model was generalizable across the folds. Further results were obtained with a training-test split of (70 *−* 30)%. The following two modalities were considered for training:

i. **Individual-Modification models**: For each of the 26 histone modifications, an individual XGBoost model was trained to predict the transcriptional class of genes and to further investigate the critical nucleosomal positions. This approach allows assessment of specific positions for each of the histone modification that contribute most significantly to predicting the class of high or low-expression genes.
ii. **Combined-Modification models**: All 26 histone modifications were concatenated to form a feature matrix of size 26 × 11 per gene, capturing positional and combinatorial information. A single XGBoost model was trained with the genes (represented as described) to predict the category of transcription. This model helps to understand how multiple histone marks jointly influence gene-expression. Additionally, to check the influence of the gene-body and promoter modifications separately (as listed below, adapted from [17]), we designed XGBoost classifiers for combinations of these specific PTMs :

- **Gene body modifications:** H3K4me, H4K20me, H3K79me3, H3K36me3, H4R3me2s, H4R3me, H3K36me, H3K36me2, H4K16ac, H3S10ph, H2AS129ph.
- **Promoter modifications:** H2AK5ac, H3K14ac, H3K18ac, H3K23ac, H3K27ac, H3K4ac, H3K4me2, H3K4me3, H3K56ac, H3K79me, H3K9ac, H4K12ac, H4K5ac, H4K8ac, Htz1.

To ensure robustness and generalizability of the models, five-fold cross-validation was performed - the dataset was divided into five equal subsets, with four subsets used for training and one subset reserved for validation (test) in each iteration. Within every fold, the data were stratified to maintain class balance and to prevent bias toward either class during model evaluation. A 70–30 training–validation split was implemented within each fold to monitor model performance and reduce the likelihood of overfitting. The rigorous cross-validation helps towards balancing bias and variance during model evaluation as recommended in previous studies [32] .

### 2.5 Hyper-Parameter Tuning

The performance of the XGBoost model was optimized through an exhaustive grid-search procedure using scikit-learn’s GridSearchCV framework. The principal hyperparameters—including the learning rate (*η*), maximum tree depth (`max_depth`), number of estimators (`n_estimators`), and regularization parameters (*λ* and *α*) were tuned to achieve optimal model performance. Multiple parameter combinations were evaluated under cross-validation to identify configurations that yielded the best balance between the F_1_-score and accuracy. Additional parameters, such as the subsampling ratio (`subsample`) and column sampling by tree (`colsample_bytree`), were adjusted to minimize overfitting and enhance model generalization, as illustrated in Fig. 1. The grid-search process ensured an appropriate trade-off between bias and variance, where the test performance guided the selection of the final model configuration. The best-performing model was then selected, and its optimal hyperparameter values were used for the experiments.

**Fig. 1.**
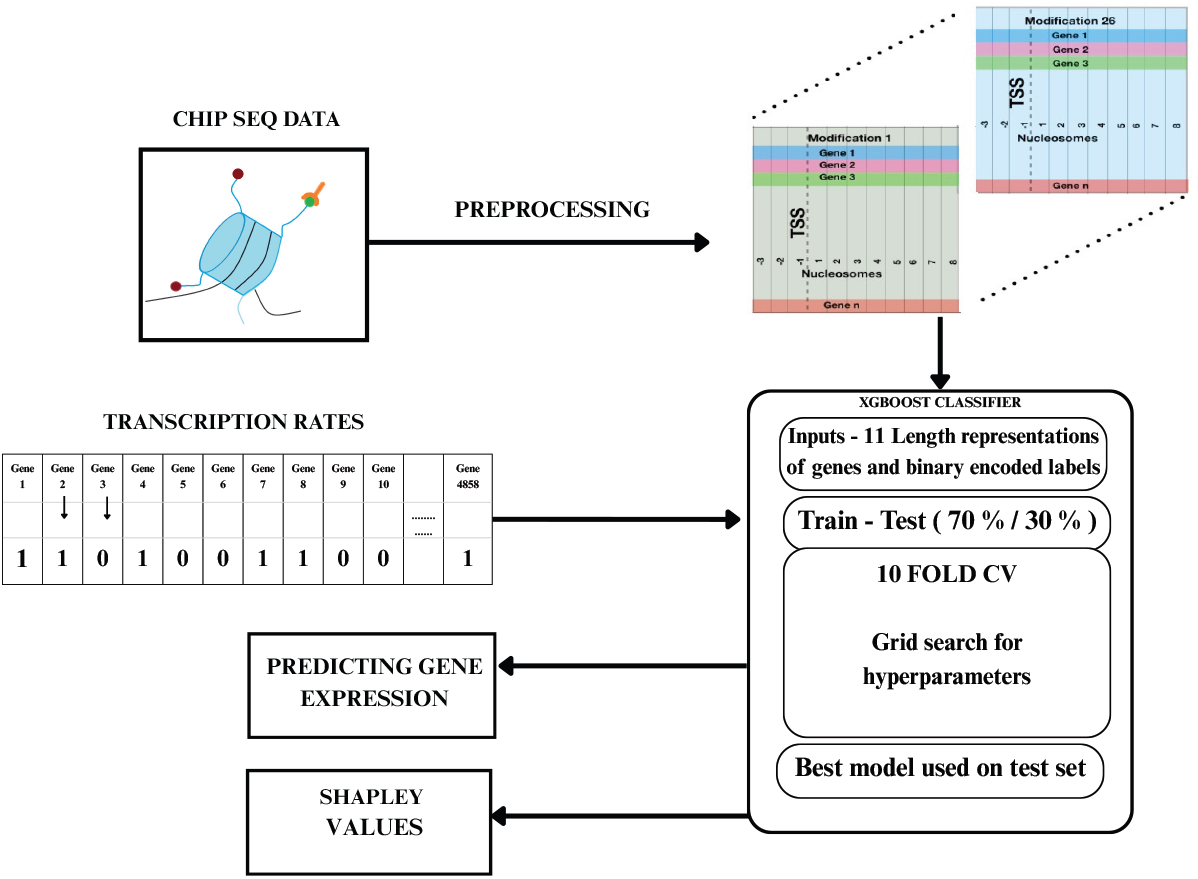
Flowchart of the methodology. ChIP-seq data containing log2 fold-change of histone PTM signals across 4800 genes; transcription rates of genes, binary encoded for classification; ChIP-seq data preprocessed into a 3D array *n*PTMs, *n*genes, *n*positions ; machine learning model (XGBoost) with hyperparameter tuning; insights drawn from the ML model.

### 2.6 SHAP (SHapley Additive exPlanations)

SHAP is a game-theoretic framework used for interpreting the predictions of any machine learning model [33]; the Shapley values quantify the contribution of each feature to a specific prediction class. In the context of the present models, SHAP was utilized to identify the most influential PTM and the specific nucleosomal positions that drive genes toward high or low expression. The python libraries *scikit − learn −* 1.8.0, *Shap −* 0.50 was employed for our study.

Analyses using the SHAP values were performed as follows:

**Gene-Level Interpretability for Individual-Modification models**: For each of the 26 individual-modification models, SHAP waterfall plots were generated for every gene to visualize how the nucleosomal positions of a histone PTM influence the predicted expression level for that gene. This approach enables the capture of gene-regulatory effects of each PTM in the context of a gene’s chromatin landscape.

- **Identification of Key Nucleosomal Positions for Promoter and Gene-body PTMs**: The positions corresponding to the highest SHAP value for every gene for the 26 histone PTM models were collected and analyzed to understand the criticality of the nucleosomal position in transcriptional regulation. This analysis was conducted separately for gene-body and promoter-associated modifications, as listed in Sec 2.4.
- **Combinatorial Patterns for the Combined-Modification model**: To study the combinatorial influence of all the histone modifications, SHAP values from the combined-modification model across the nucleosomal positions for each gene were analyzed. By obtaining SHAP importance scores across PTMs in the nucleosomal positions, we studied which modification and its respective position might be the key players together towards gene regulation. This analysis enables the understanding of modification patterns that work synergistically or antagonistically with each other during transcription activity - to validate the histone-code hypothesis.

Algorithm 1 outlines the workflow for classifying gene transcription rates using histone PTM signals across nucleosomal positions. It includes data normalization, median-based label assignment, feature construction, and cross-validated evaluation of multiple machine-learning models.

## 3 RESULTS

### 3.1 Active and inactive genes are predicted using the patterns of histone PTMs

The XGBoost models for the individual-modifications and the combined-modification were evaluated based on their F-1 scores and accuracies towards prediction of the transcriptional rate of the gene - Fig. 2 shows these values for all the PTMs and the combined model. It can be inferred from this figure that the combined-modification model (with an accuracy and F1 score of 70% and 83% respectively) outperforms the individual-modification models. The F1 and accuracy scores for the combined gene-body and promoter PTMs were higher than most of the individual PTMs, but were lower than the combined-modification model - thereby establishing both the sets (gene-body and promoter) modifications are required for determining the transcription rate of the gene.

**Fig. 2.**
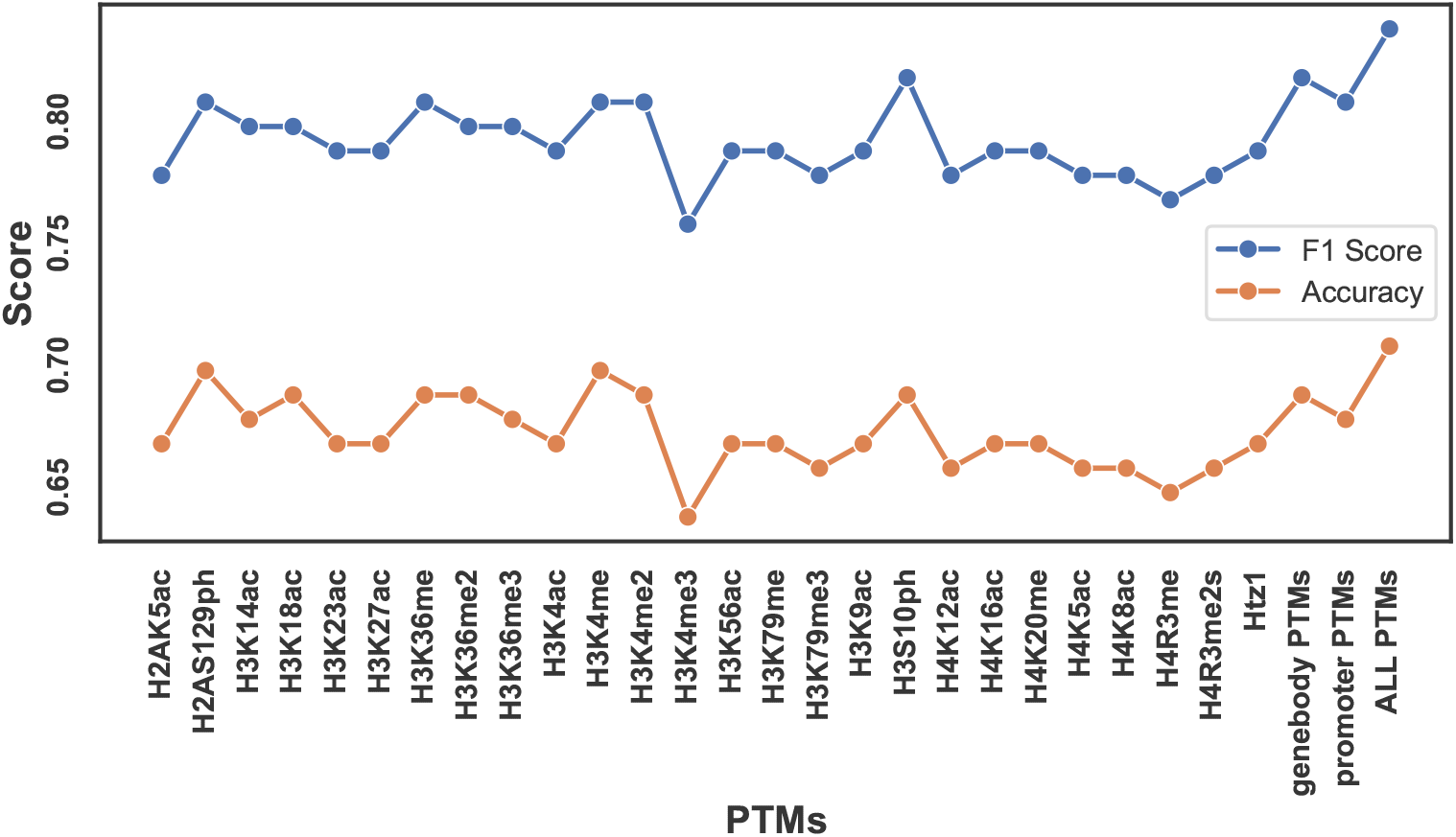
Performance of the XGBoost model across individual histone modifications. The line plot compares F1-scores (circles) and accuracies (squares) of the XGBoost classifier when trained on individual histone modifications. Specific modifications exhibit higher metrics, with F1-scores exceeding 80∥ for several cases. These results emphasize that not all histone modifications contribute equally to transcriptional regulation, and a subset of them play a strong role in influencing gene-expression states.

#### Algorithm 1

Gene-transcription rate classification from histone PTMs

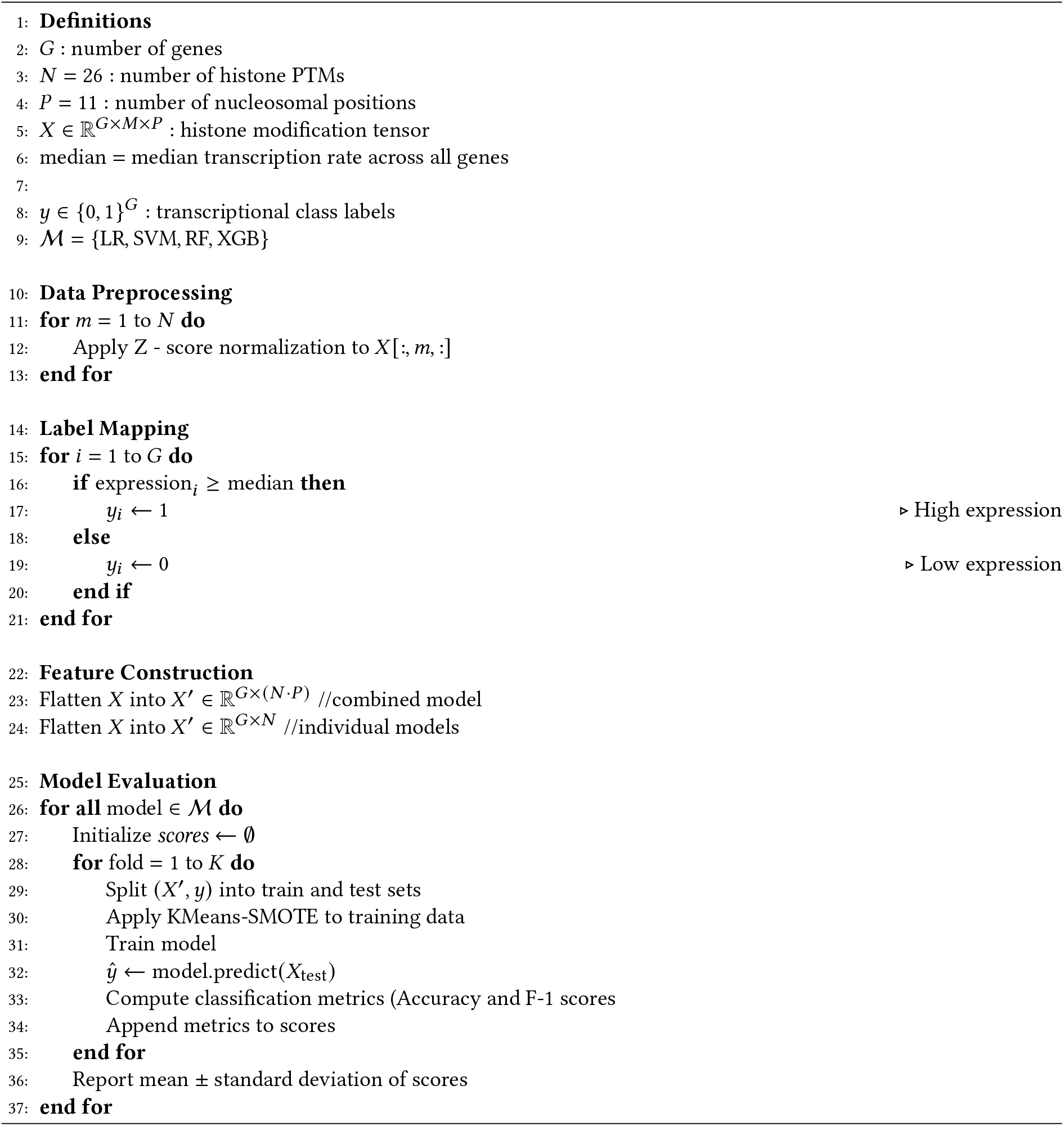

Individual-modification models from PTMs such as H3S10ph, H3K4me2 and H3K4me provide higher scores while H3K4me3 provided the least accuracy and F1 scores of 63% and 75% respectively. It should also be noted that the gene-body PTMs or the promoter modifications do not show any distinction in terms of the resultant scores, when applied individually.

We now analyze the waterfall plots for the XGBoost models based on each of the 26 PTMs for the genes - these plots are particularly informative as they provide a gene-specific understanding of how each of the nucleosomal positions influence the prediction of gene-expression for the specific histone PTM. A clear distinction was observed in the waterfall plots between the transcriptionally active and repressed genes: the SHAP values of the active genes (e.g. AAC1) were dominated by the positive SHAP coefficients, while the negative coefficients dominated the prediction of a repressed gene such as AAH1 - see Fig. 3 for two PTMs (H3K36me3 and H3K27ac). The waterfall plots for these genes for the other PTMs are provided in SI (Fig. S2-S3). It has to be noted that the most influential positions for the genes vary across the PTMs - which we analyze in the next subsection.

**Fig. 3.**
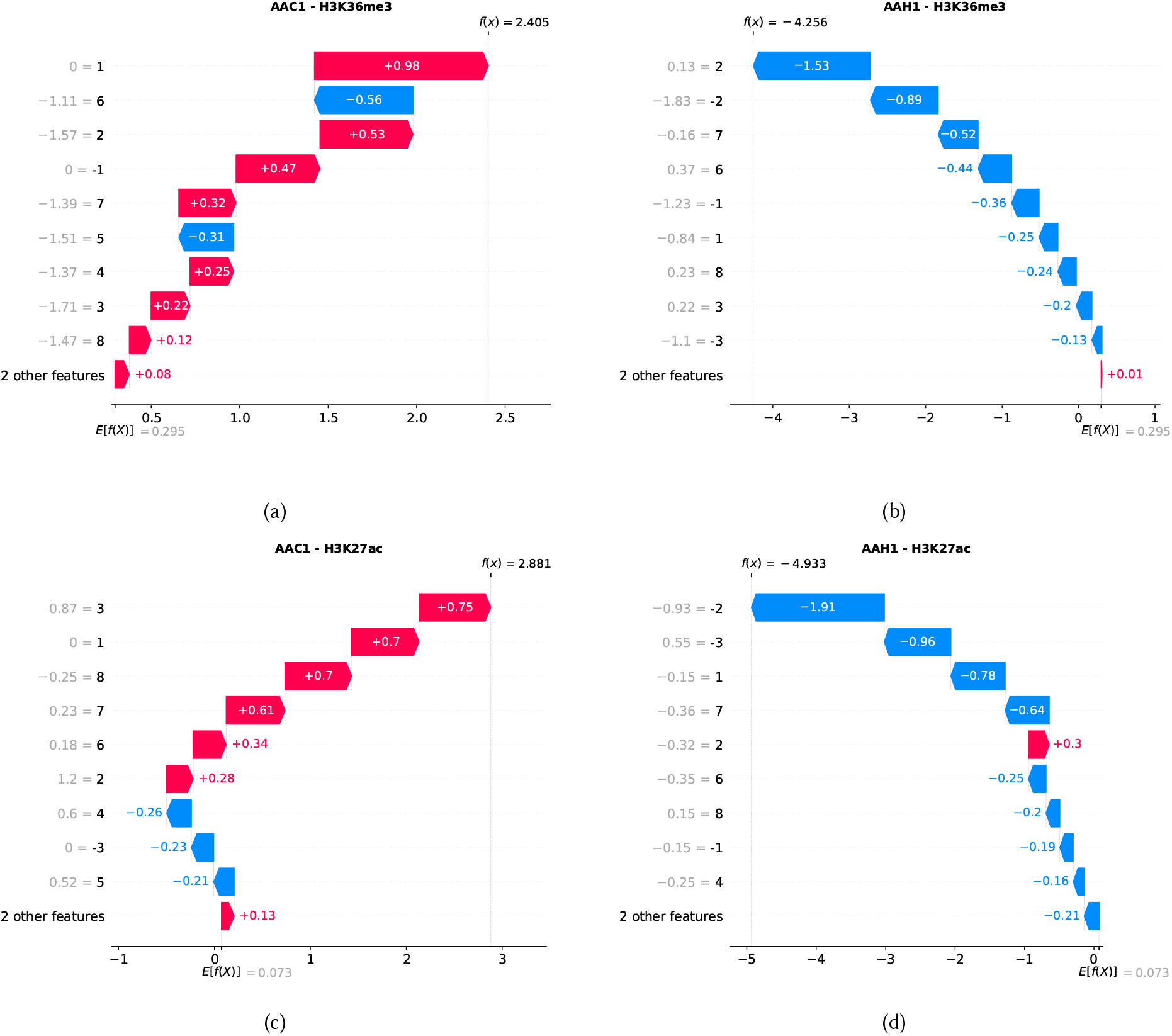
(a-d) SHAP-based positional importance plots for two histone modifications (H3K36me3 and H3K27ac) for two representative genes: AAC1 (high-expression) and AAH1 (low-expression).

### 3.2 Distinct nucleosomal positions critically influence various PTMs

To further interpret the critical influence at specific nucleosomal positions, a metric corresponding to the probability of a position being the most critical is defined for every PTM as follows : the number of genes for which that position had the highest absolute SHAP relative to all the correctly predicted genes. The resulting probabilities of criticality of the positions for each PTM are plotted in Fig. 4a. It can be inferred that most positions have about (5 *−* 12)% probability of being the top influencers for some (but not all) PTMs while position 2 has a higher probability of being the most influential position for 3 PTMs - H4K5ac, H3K4me and H3K14ac. Positions 1, 5 and 7 are also key locations in the gene body for certain PTMs, influencing the gene-expression.

**Fig. 4.**
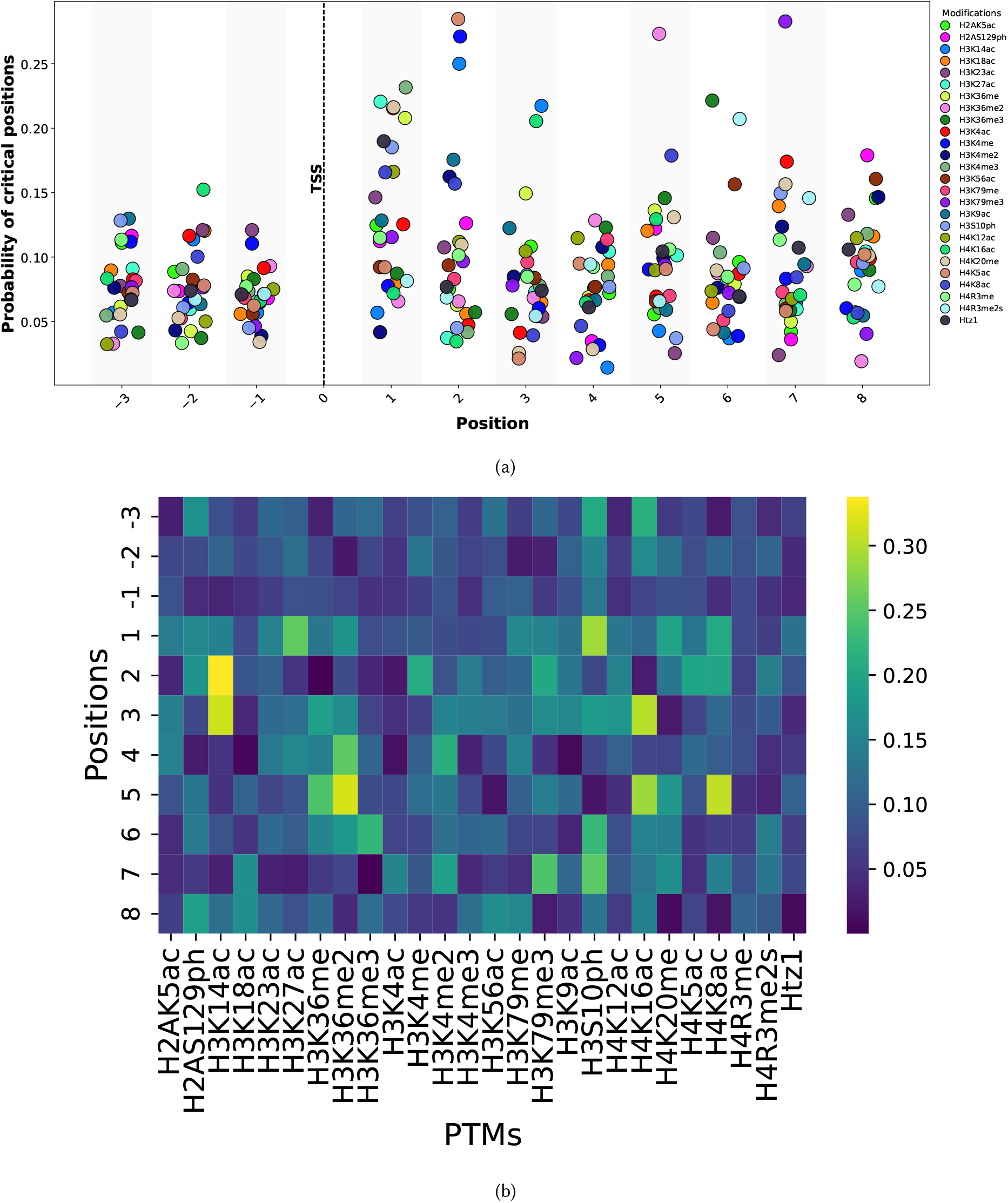
SHAP-based positional importance of PTMs . (a) Plot showing the probability distribution of the most influential nucleosomal positions based on SHAP values, combining all histone PTMs into a single figure. The probabilities of critical positions are computed as describsed in subsection 3.2. (b) Heatmap showing the average SHAP values for each position across the 26 PTMs.

To further understand the contribution of PTMs, we generated a heatmap of the SHAP values of each of the 11 positions for the 26 histone modifications, averaged over all the genes. The results from this heatmap corroborated with probabilities of criticality from Fig 4b.

For H3K14ac, the highest average SHAP value was observed at position +2, illustrating its strong role in early gene body regulation - it should be noted that position 2 showed a very high probability of being the most critical position for H3K14ac from Fig. 4a. For H3K36me2, the highest average SHAP was observed at position +5, showing strong influence in transcribed regions. These findings underline the hypothesis that both the type of modification and its presence in specific positions are critical in understanding transcriptional activity. We also obtained stacked bar plots for the average SHAP values for the active and inactive genes separately at each position (Fig. S4 and S5 from SI for gene-body and promoter PTM models), where blue (orange) bars denote the average SHAP values of the histone PTM at that position for the correctly predicted active (inactive). The presence of both the positive and negative contributions across the positions reflect how activating and repressive marks exert distinct, position-specific effects on transcriptional regulation for different genes.

Additionally, the magnitude of the average SHAP value reflects the overall importance of a given histone modification at a specific nucleosomal site, thereby highlighting which positions are most influential for gene-expression prediction. For instance, in case of PTM H3K14ac (Fig. S6(b) from SI), while position *−*1 does not have any influence towards prediction of active genes, it has a significant value towards prediction of inactive genes - it is interesting to note that the opposite effect is exerted by position 1 for the same histone PTM. The beeswarm plots for the individual models ( where each dot corresponds to an individual gene, with its location along the x-axis indicating the direction and magnitude of the contribution toward high or low expression) are also plotted in Fig. S6 and S7 of the SI.

### 3.3 Combinatorial interactions between histone modifications and nucleosomal positions drive gene-expression prediction

To understand the combinatorial influence of all the 26 histone PTMs and their positional signals on gene-expression, we trained an XGBoost model with an input vector dimension (26 × 11) for each gene (as described in 2.4 for the combined model). This model achieved an F1-score of 82% and an accuracy of 70%, demonstrating how the interplay between these PTMs is pivotal for transcriptional regulation. The beeswarm plot of Fig. 5a highlights the top 20 most influential PTM - position pair influencing gene-expression prediction. From the beeswarm plot, one can observe that the activating PTMs such acetylations on H3 (and their gene-body positions) are amongst the top influencers of gene-expression.

**Fig. 5.**
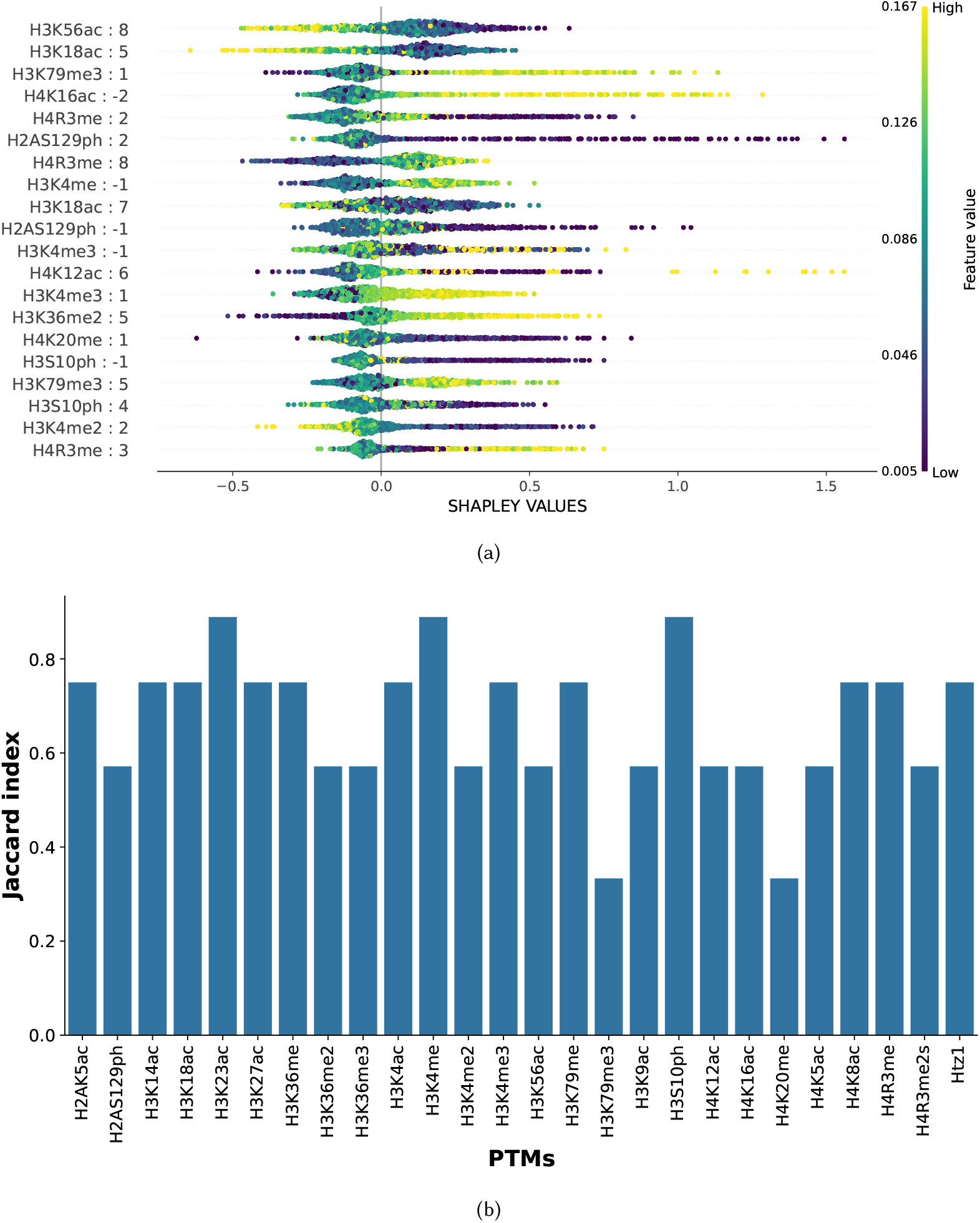
(a) The SHAP beeswarm plot shows uneven feature importance, with a small subset of modification–position pairs exerting dominant influence on gene expression. (b) Jaccard index values for different PTMs, highlighting the similarity of top-5 PTMs across individual and combined models.

To see the consistency between important features of the individual-PTM models and the combined-PTM model, we computed the Jaccard similarity index [34] between the sets of top five positions from each individual histone PTM model and those from the same PTM extracted from the combined modification model. From the results plotted in Fig. 5b, the higher values of Jaccard similarity indices indicate a significant overlap between the two types of models, indicating that the same positions are critical in the individual models and the combined model. Certain PTMs such as H3K79me3 and H4K20me have Jaccard similarity indices less than 0.5 suggesting that the positional importance of the individual and combined models are different.

## 4 DISCUSSION

In this study, we employ ML models coupled with SHAP explainability to decipher critical PTMs and their positions along the genes for transcriptional regulation in yeast. Initially, we represent the 26 histone PTM dataset in the steady-state yeast study [17], as 11 positions (3 positions in the promoter and 8 positions in the gene-body) for each gene. This was followed by application of ML models trained on the PTM datasets to predict the transcription rate of genes, discretized as fast (≥ 13 mRNA/hr) or slow (*<* 13 mRNA/hr). While several ML models such as Logistic Regression, SVM and Random Forests were explored (results in the SI - Fig.S1), the XGBoost model gave reasonably high values of accuracy and F1 scores for all the 26 PTMs, combined gene-body, combined promoter and all the PTM based models.

In the era of XAI models, where interpretability of Machine Learning is critical, we have utilized the SHAP explainability metrics to decipher the positional importance of the histone PTMs amidst their cross-talk and their patterns across the genes in yeast. Several similar groups have attempted similar studies towards interpreting different regulatory mechanisms in the past Karlić et al. showed that gene expression in yeast can be accurately modeled using histone modification signals, providing proof for a chromatin-based regulatory code [18]. However, their analysis relied on linear models and region-averaged PTM signals, which limited the ability to look for effects along genes or capture non-linear interactions between multiple modifications and how these modification differ in their activity across various positions along the gene. Weiner et al. generated genome-wide maps of 26 histone modifications in yeast under steady-state and stress conditions, showing how different modification patterns at promoters and gene bodies are associated with transcriptional activity [17]. This study provided a detailed descriptive view of where histone PTMs are located and how they change with gene activity, but it did not quantitatively assess how much each modification or each position contributes to transcriptional output. In our work, we build directly on this dataset by applying supervised machine-learning models and using SHAP explainability to interpret their predictions. SHAP allows us to assign importance scores to individual histone modifications at specific promoter and gene-body positions, making it possible to identify which PTMs, and which locations along the gene, most strongly influence transcriptional states. This approach moves beyond descriptive mapping and enables a more interpretable and position-resolved understanding of how histone modifications collectively regulate gene expression in yeast.

While most of the individual PTM based models gave reasonably high accuracy and F1-scores, the combined models ((i)with only gene-body PTMs, (ii) with only promoter PTMs, (iii) with all PTMs ) outperformed the individual models. The same positions (such as 2) had positive coefficients of SHAP, for active genes (eg.AAC1) and negative coefficients for inactive) genes (such as AAH1) for the same PTM (such as H3K36me3). Another important inference from our study is that certain positions such as 2, 5, 7 for certain specific PTMs such as H3K14ac, H3K36me2, H4K5ac, H3K79me3 had high average SHAP values indicating their important roles in the prediction of many genes.

The beeswarm SHAP plots quantified the importance of individual PTMs (and their positions as a pair) by ranking them in the order of importance. Activating PTMs such as H3K56ac (position 8), H3K18ac (postion 5), H3K79me3 (position 1) were identified as the highest contributing features across the genes towards their transcriptional activity prediction. The Jaccard similarity indices were obtained for every PTM (across a set of top 5 positions in the individual PTM model and another set of top 5 positions in the combined model for that PTM) and analysed - the high values indicated that the high SHAP positions were common across the individual and combined PTM models.

The shortcomings of our approaches include the imbalance in the datasets (with more genes being transcriptionally active than inactive) which we have circumvented through the use of KMeans-SMOTE for training the models. Additionally the presence of false-positives and false-negatives indicate that the models may not be fully generalizable; however, these incorrectly predicted genes were not considered for SHAP analysis. In the future, we intend to employ Deep Learning models such as Transformers that understand the spatial context between the modifications and their explainability tools for interpretation across more genes.

While the current study focuses on the inter-relationships between the PTMs and their positional significance across the genes, considering the DNA methylation patterns and related epigenetic signals in gene regulation would aid in better prediction - which is considered as future work. We would also aim to extend this study to other model organisms to study the evolutionary variations of the SHAP scores of the PTMs and their patterns, as subsequent work.

