## Supplementary material for "An Explainable Machine Learning Approach to study the positional significance of histone post-translational modifications in gene regulation": SI

Sharun Ramachandran Nithya Ramakrishnan

### Supplementary Information

#### 1.1 Comparison of Model Performance on Histone Modification Features

Comparison of F1 scores across histone modifications (PTMs) for all machine learning models — Random Forest (RF), XGBoost, SVM (Support Vector Machines) , and Logistic Regression. Although the Random Forest model achieved the highest overall F1 score, it exhibited signs of overfitting and predicting only one class hence the high f1 score and was therefore excluded from SHAP-based interpretability analysis. XGBoost showed more consistent generalization performance across PTMs

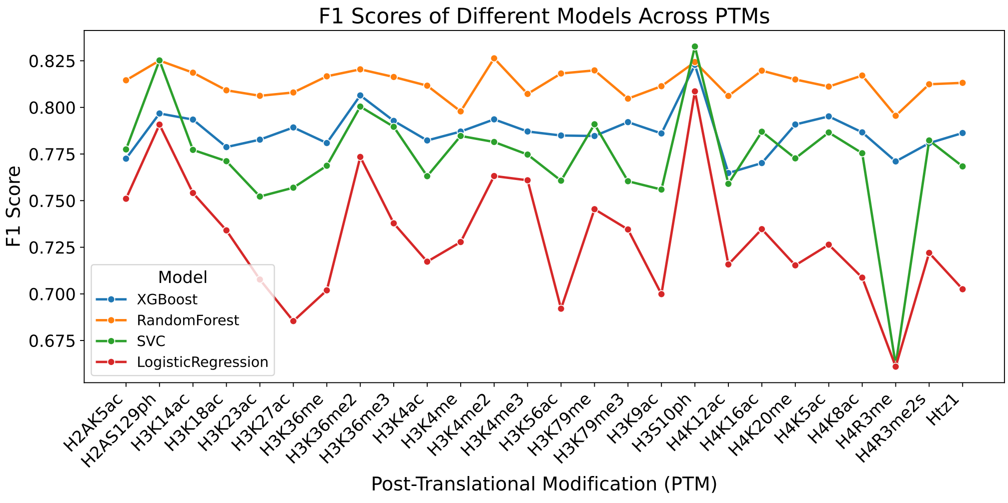


Fig. S1 Plot showing the F1 scores of three machine learning models — Random Forest (RF), XGBoost, and Logistic Regression — across different histone modifications (PTMs). Each line represents one model, with distinct colors indicating model type. The x-axis lists the PTMs, and the y-axis denotes the corresponding F1 score.

#### 1.2 Interpreting how each nucleosomal position and PTM pushes a gene toward high or low expression through waterfall plots

To understand of the contribution of PTMs and positional features to gene expression classification, we employed SHAP (SHapley Additive exPlanations) waterfall plots. These plots provide

| Modification | Acc. Mean | Acc. Std | F1 Mean | F1 Std |
|---|---|---|---|---|
| H2AK5ac | 0.653 | 0.016 | 0.774 | 0.012 |
| H2AS129ph | 0.680 | 0.015 | 0.799 | 0.011 |
| H3K14ac | 0.663 | 0.020 | 0.782 | 0.015 |
| H3K18ac | 0.667 | 0.019 | 0.786 | 0.014 |
| H3K23ac | 0.654 | 0.019 | 0.773 | 0.014 |
| H3K27ac | 0.644 | 0.026 | 0.764 | 0.019 |
| H3K36me | 0.669 | 0.017 | 0.789 | 0.012 |
| H3K36me2 | 0.670 | 0.021 | 0.789 | 0.015 |
| H3K36me3 | 0.653 | 0.025 | 0.774 | 0.018 |
| H3K4ac | 0.655 | 0.020 | 0.775 | 0.014 |
| H3K4me | 0.669 | 0.018 | 0.790 | 0.013 |
| H3K4me2 | 0.676 | 0.023 | 0.793 | 0.019 |
| H3K4me3 | 0.641 | 0.021 | 0.765 | 0.016 |
| H3K56ac | 0.673 | 0.012 | 0.791 | 0.010 |
| H3K79me | 0.659 | 0.019 | 0.781 | 0.013 |
| H3K79me3 | 0.657 | 0.021 | 0.776 | 0.017 |
| H3K9ac | 0.655 | 0.016 | 0.774 | 0.012 |
| H3S10ph | 0.689 | 0.016 | 0.809 | 0.011 |
| H4K12ac | 0.634 | 0.014 | 0.760 | 0.011 |
| H4K16ac | 0.660 | 0.021 | 0.782 | 0.017 |
| H4K20me | 0.665 | 0.023 | 0.785 | 0.017 |
| H4K5ac | 0.648 | 0.027 | 0.771 | 0.021 |
| H4K8ac | 0.649 | 0.020 | 0.772 | 0.017 |
| H4R3me | 0.631 | 0.022 | 0.755 | 0.018 |
| H4R3me2s | 0.640 | 0.025 | 0.764 | 0.019 |
| Htz1 | 0.647 | 0.016 | 0.770 | 0.011 |

Table S1. Performance of individual histone modification models obtained using 10-fold cross-validation.

a local explanation of the model's decision for a single gene by breaking down the prediction into the additive contributions of individual features relative to a baseline value.

In each plot, the baseline value corresponds to the model's expected output across the training dataset. Starting from this baseline, features sequentially "push" the prediction higher or lower depending on whether their SHAP value is positive or negative. Features with positive contributions (shown as rightward bars,RED) increase the probability of the gene being classified into the high-expression group, while features with negative contributions (leftward bars,BUE) push the prediction towards the low-expression class. The final prediction is obtained by summing the baseline and all feature contributions.

#### 1.3 Gene - AAC1

The figures presents SHAP (SHapley Additive exPlanations) waterfall plots showing the contribution of 26 histone modifications to the high expression prediction of the AAC1 gene. Each plot represents one histone modification, showing how individual positional features within that modification influence the model's output toward the "high expression" (+1) class.

Positive SHAP values (red bars) indicate features that increase the model's confidence in the high-expression prediction, whereas negative SHAP values (blue bars) represent features that suppress the prediction. The base value denotes the model's expected output before looking at any of the feature which in our case are positions, and the final SHAP expectation value show how the combination of positional modification signal values leads to the final model prediction.

Overall, the waterfall plots show the importance and influence of each histone modification across nucleosomic positions for AAC1. These insights provide interpretability of how the trained model relates nucleosomic positions with gene expression regulation .

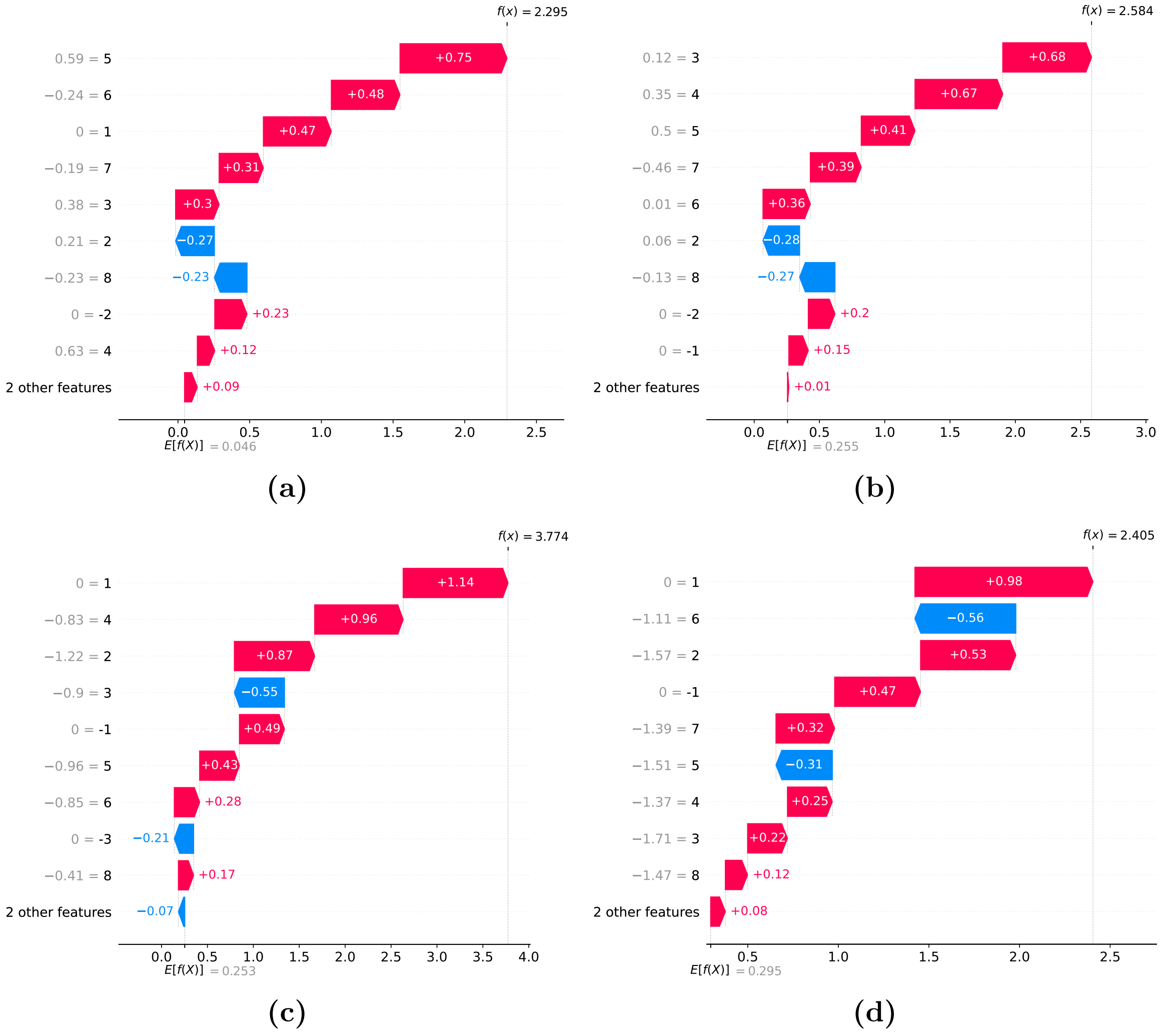


Fig. S2: (a–d: H3K4me, H4K20me, H3K79me3, H3K36me3) SHAP waterfall plots for representative histone modifications — yeast gene *AAC1*.

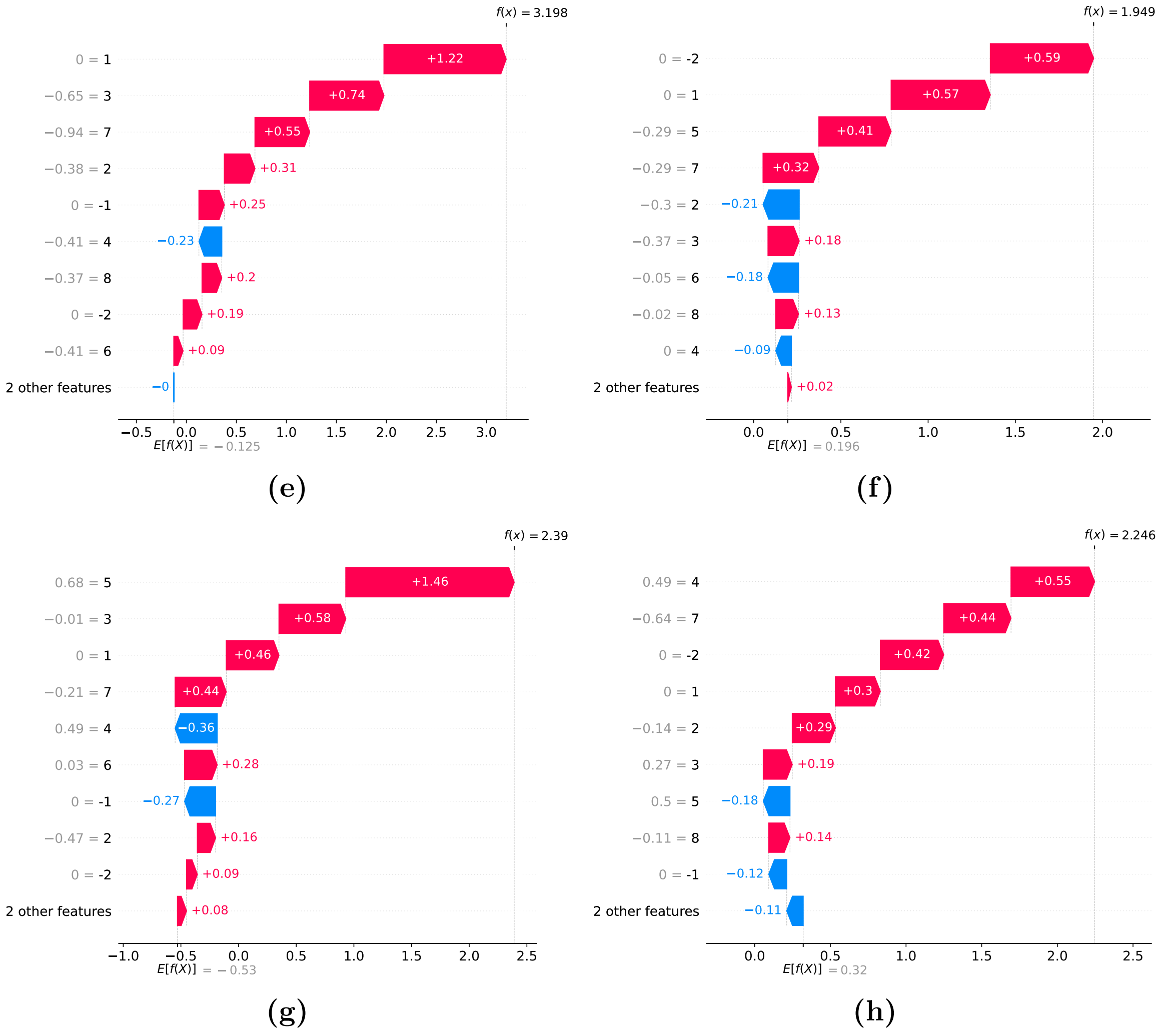


Fig. S2: (e–h: H4R3me2s, H4R3me, H3K36me, H3K36me2) SHAP waterfall plots for histone modifications — yeast gene *AAC1*.

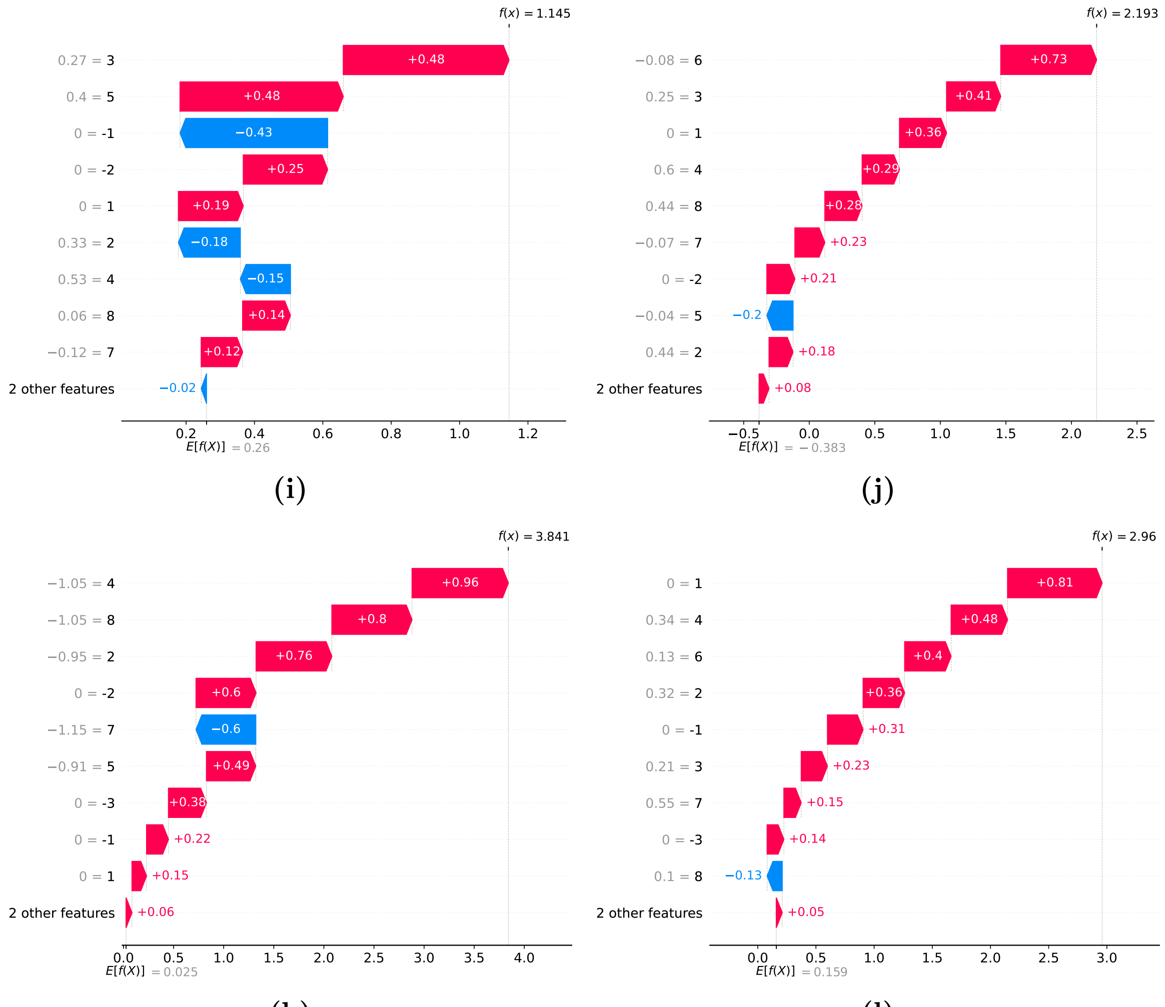


Fig. S2: (i–l: H4K16ac, H3S10ph, H2AS129ph, H2AK5ac) SHAP waterfall plots for histone modifications — yeast gene *AAC1*.

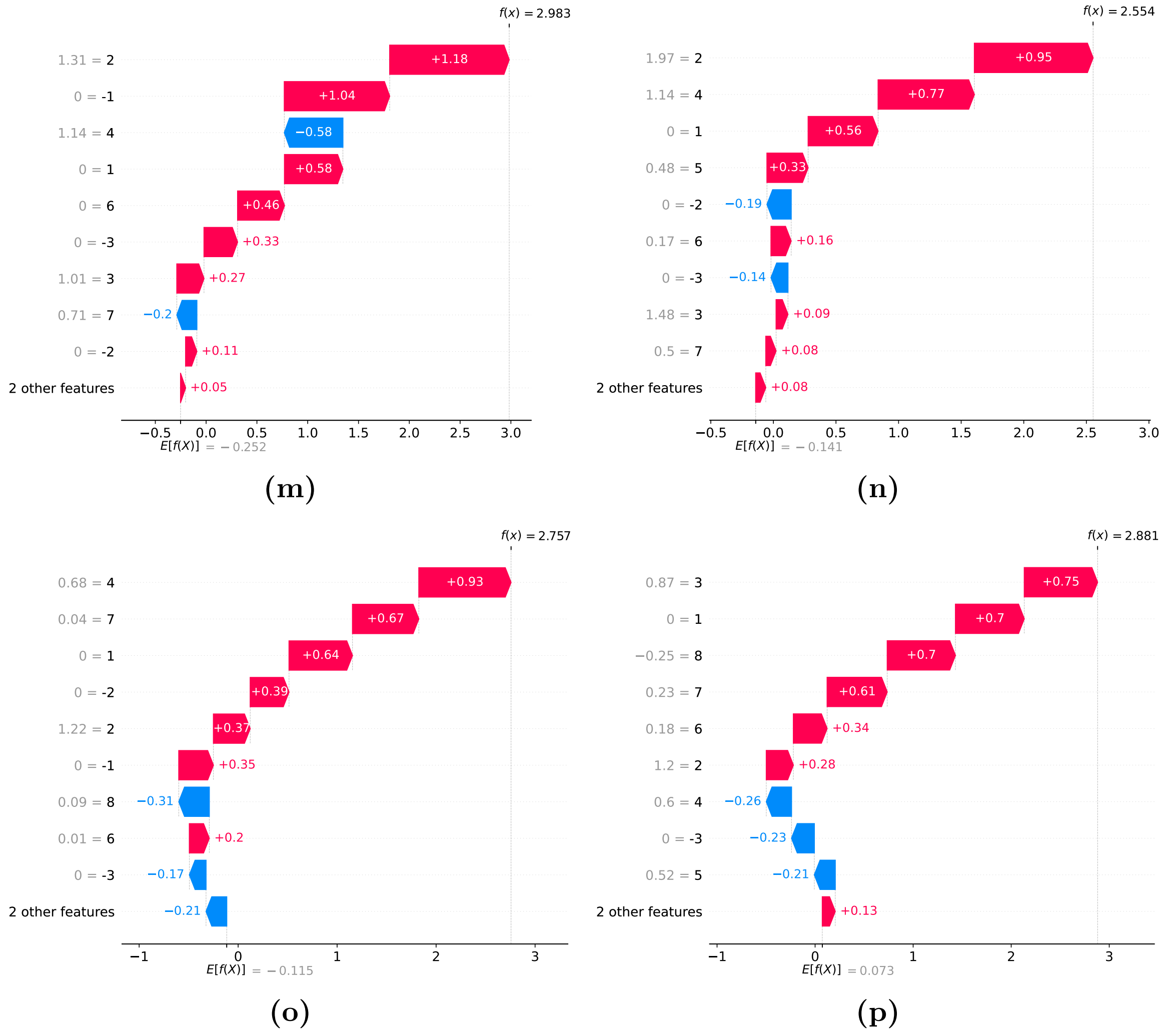


Fig. S2: (m–p: H3K14ac, H3K18ac, H3K23ac, H3K27ac) SHAP waterfall plots for histone modifications — yeast gene *AAC1*.

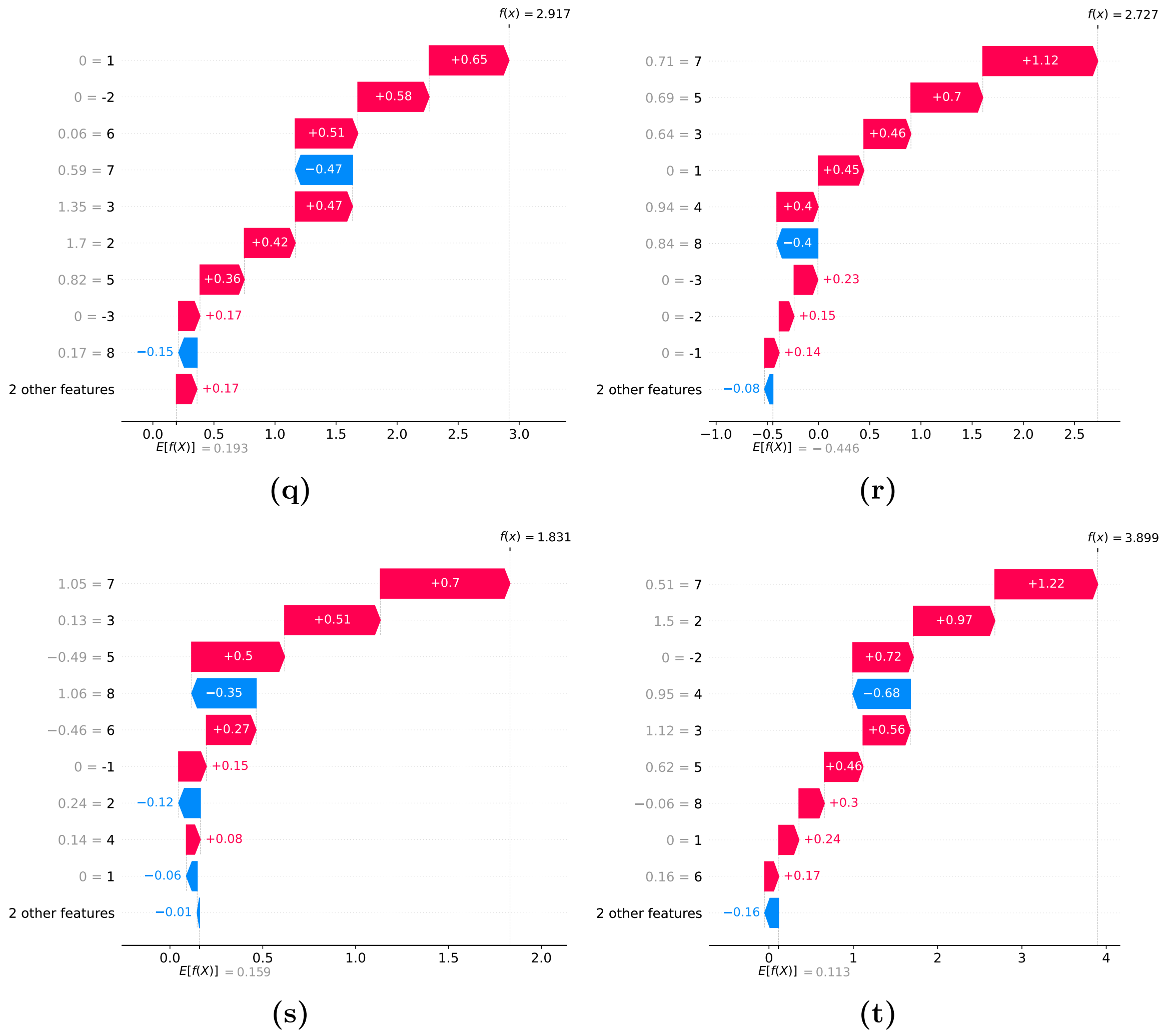


Fig. S2: (q–t: H3K4ac, H3K4me2, H3K4me3, H3K56ac) SHAP waterfall plots for histone modifications — yeast gene *AAC1*.

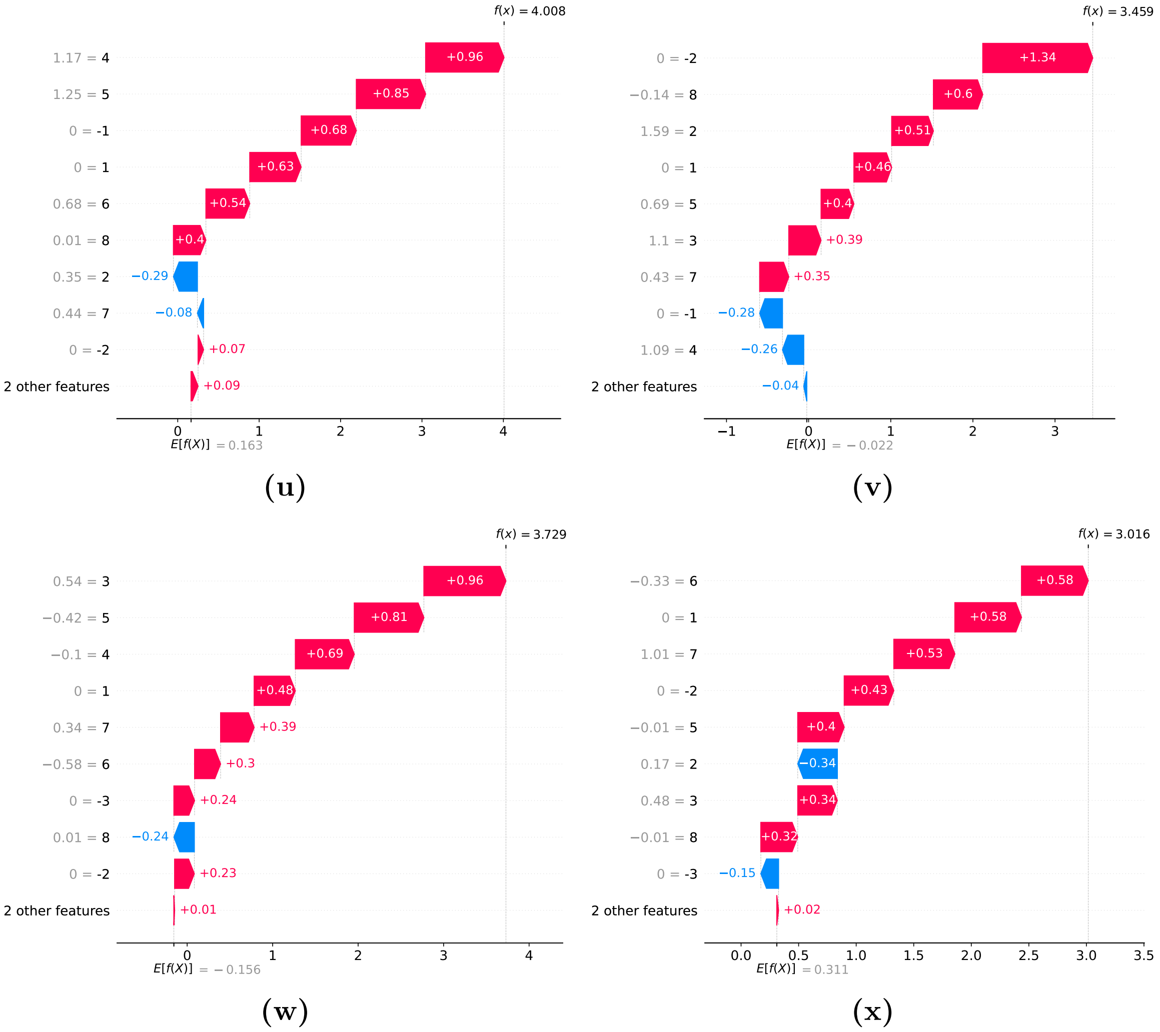


Fig. S2: (u–x: H3K79me, H3K9ac, H4K12ac, H4K5ac) SHAP waterfall plots for histone modifications — yeast gene *AAC1*.

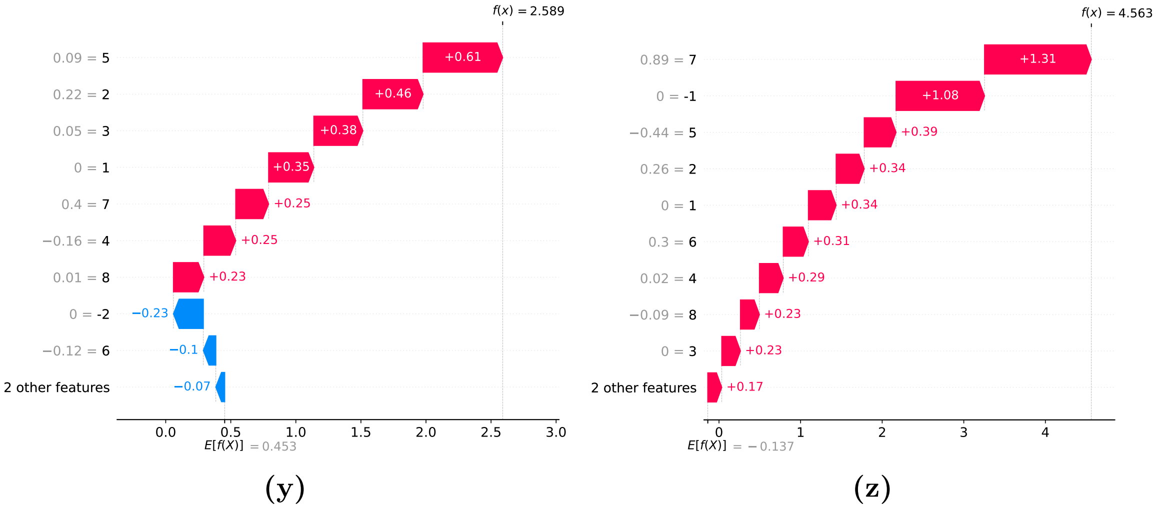


Fig. S2: (y–z: H4K8ac, Htz1) SHAP waterfall plots for histone modifications — yeast gene *AAC1*.

### 1.4 AAH1

The figures presents SHAP (SHapley Additive exPlanations) waterfall plots showing the contribution of 26 histone modifications to the low expression prediction of the AAH1 gene. Each plot represents one histone modification, showing how individual positional features within that modification influence the model's output toward the "low expression" (0) class.

Positive SHAP values (red bars) indicate features that increase the model's confidence in the high-expression prediction, whereas negative SHAP values (blue bars) represent features that suppress the prediction. The base value denotes the model's expected output before looking at any of the feature which in our case are positions, and the final SHAP expectation value show how the combination of positional modification signal values leads to the final model prediction.

Overall, the waterfall plots show the importance and influence of each histone modification across nucleosomic positions for AAH1. These insights provide interpretability of how the trained model relates nucleosomic positions with gene expression regulation .

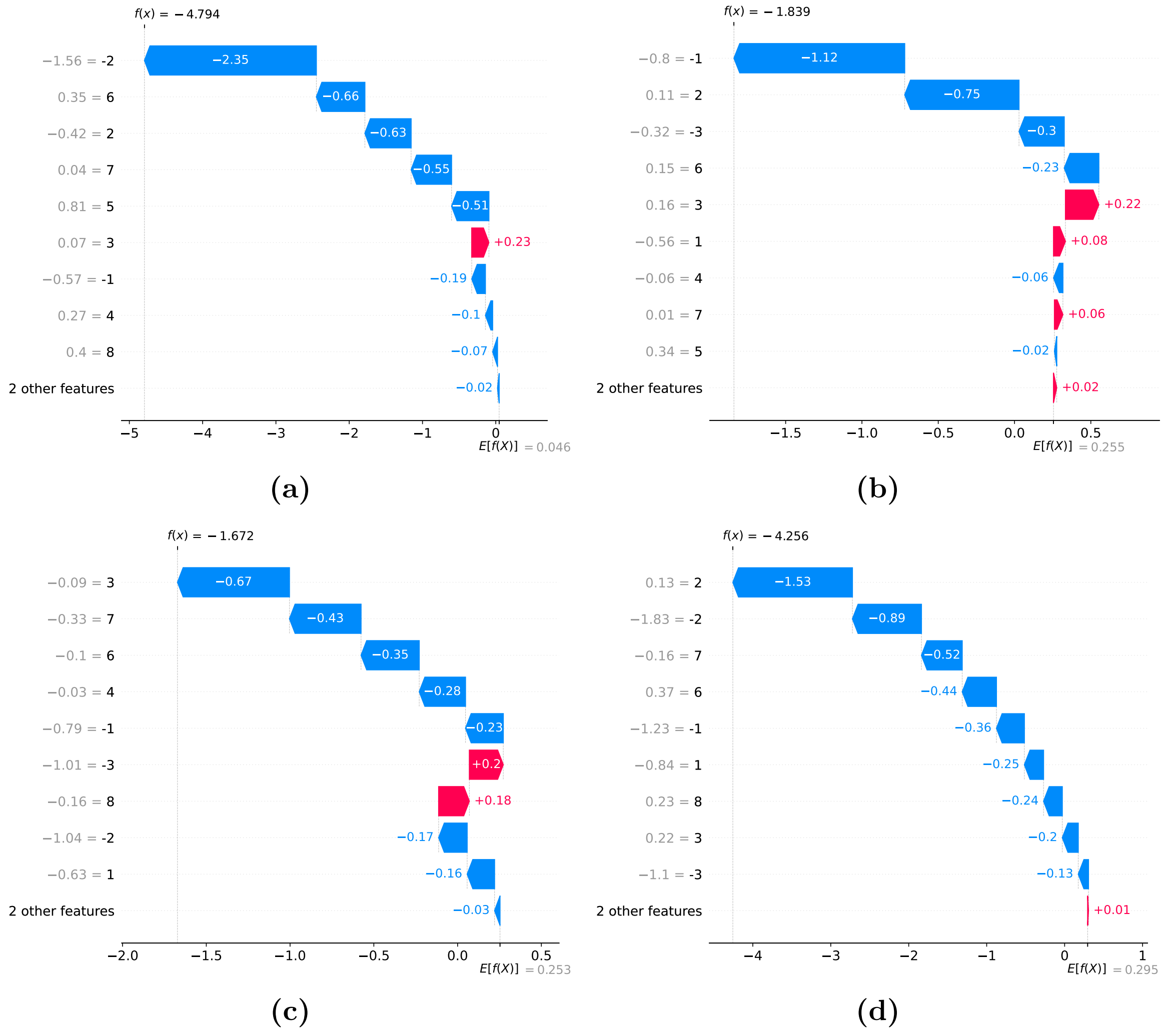


Fig. S3: (a–d: H3K4me, H4K20me, H3K79me3, H3K36me3) SHAP waterfall plots for representative histone modifications — yeast gene *AAH1*.

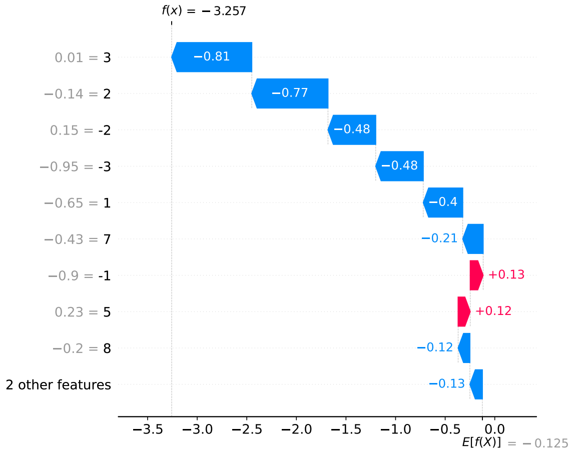


**(e)**

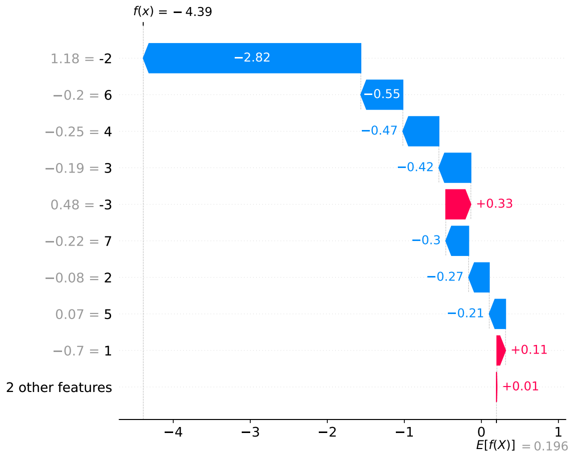


**(f)**

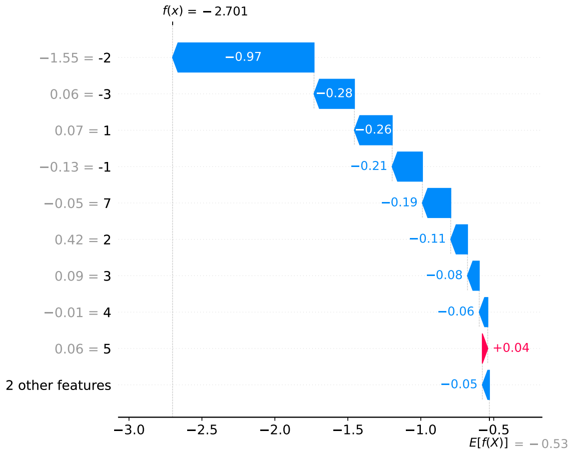


**(g)**

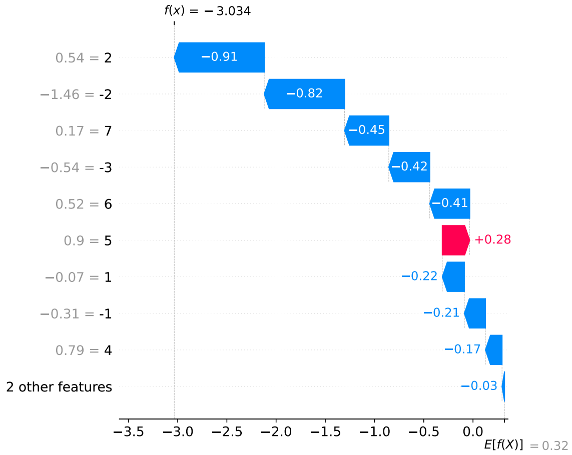


**(h)**

Fig. S3: (e–h: H4R3me2s, H4R3me, H3K36me, H3K36me2) SHAP waterfall plots for histone modifications — yeast gene *AAH1*.

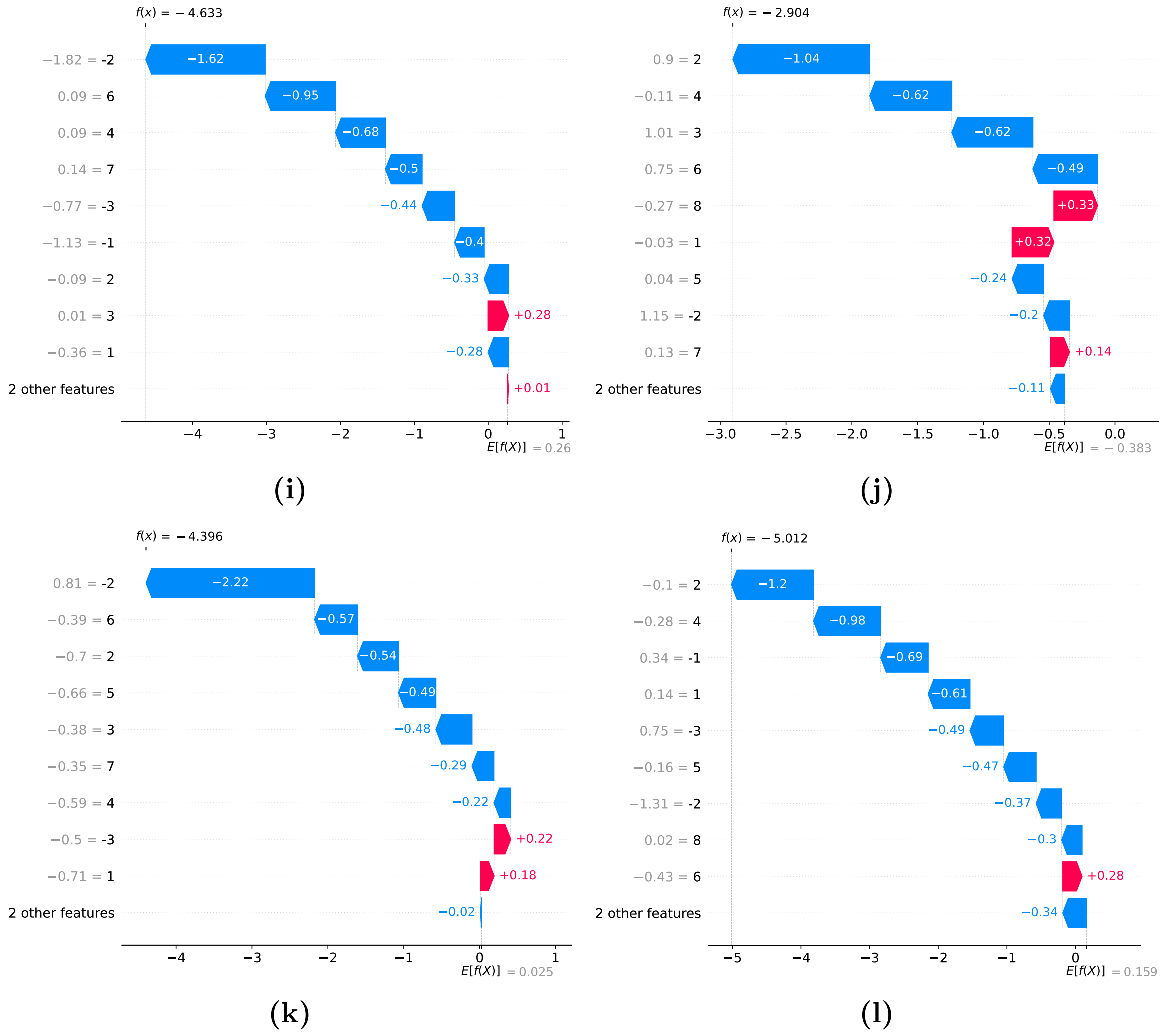


Fig. S3: (i–l: H4K16ac, H3S10ph, H2AS129ph, H2AK5ac) SHAP waterfall plots for histone modifications — yeast gene *AAH1*.

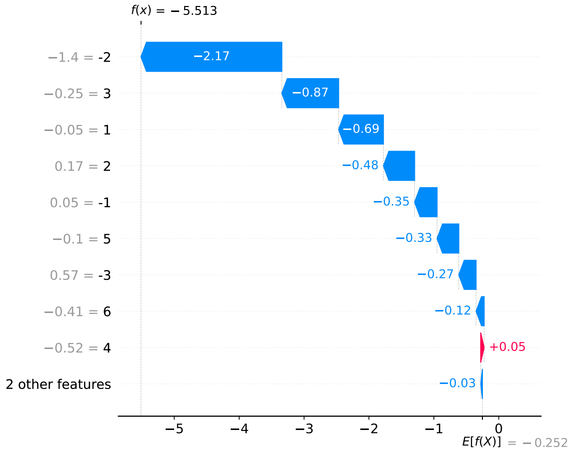


**(m)**

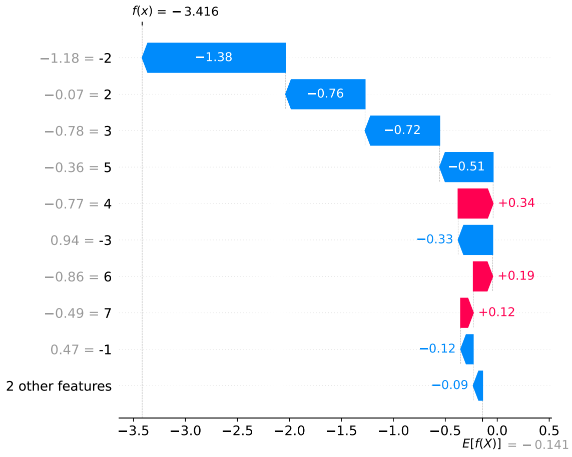


**(n)**

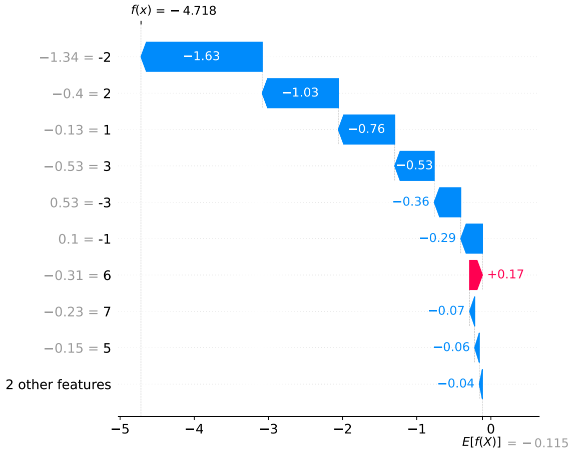


**(o)**

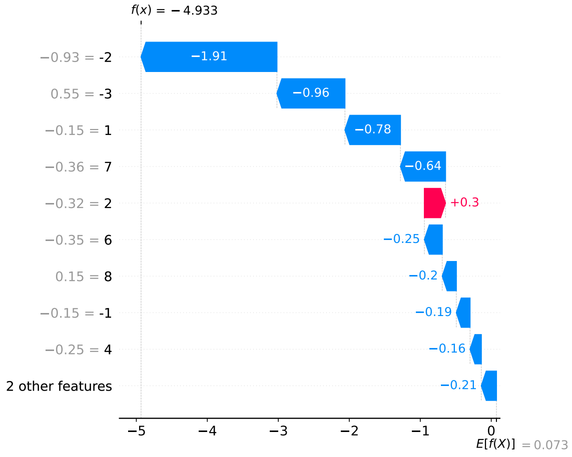


**(p)**

Fig. S3: (m–p: H3K14ac, H3K18ac, H3K23ac, H3K27ac) SHAP waterfall plots for histone modifications — yeast gene *AAH1*.

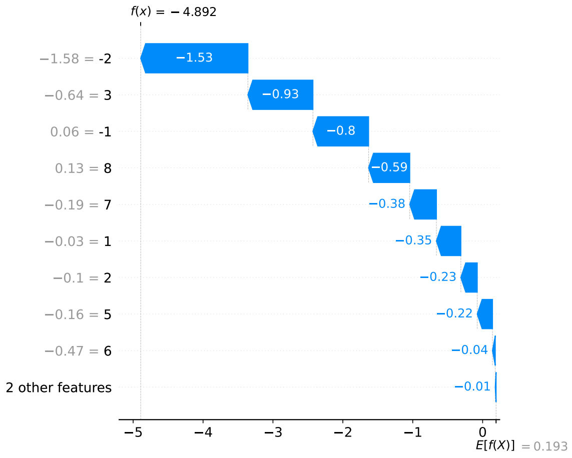


(q)

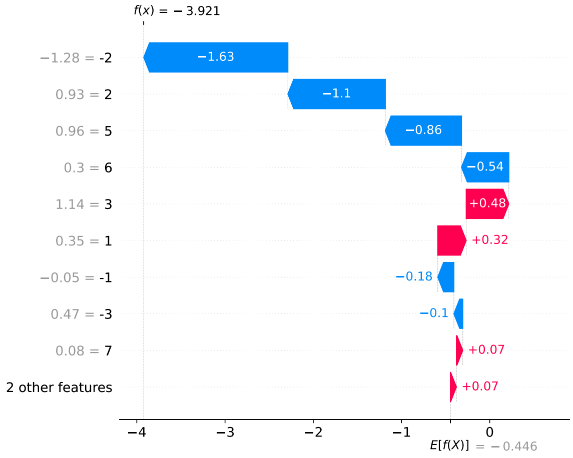


(r)

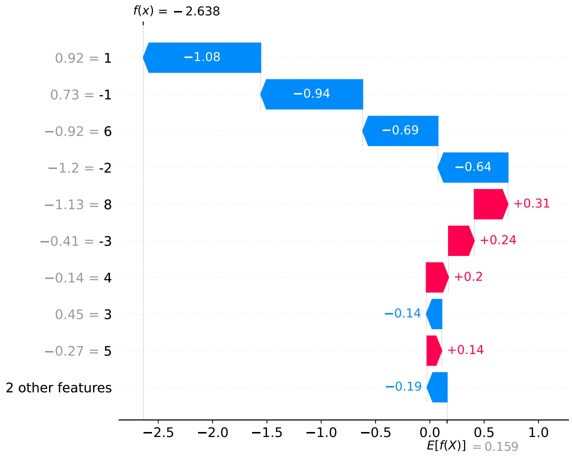


(s)

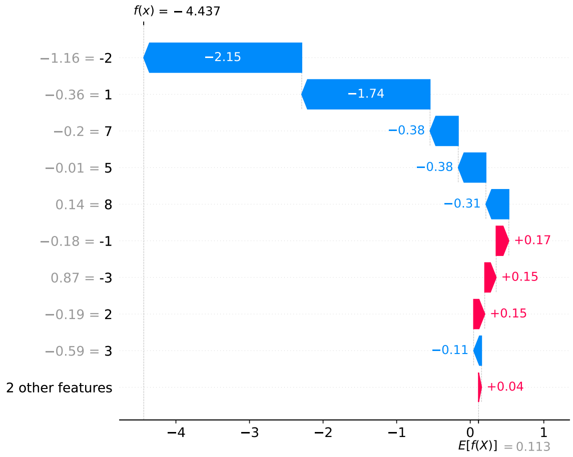


(t)

Fig. S3: (q–t: H3K4ac, H3K4me2, H3K4me3, H3K56ac) SHAP waterfall plots for histone modifications — yeast gene *AAH1*.

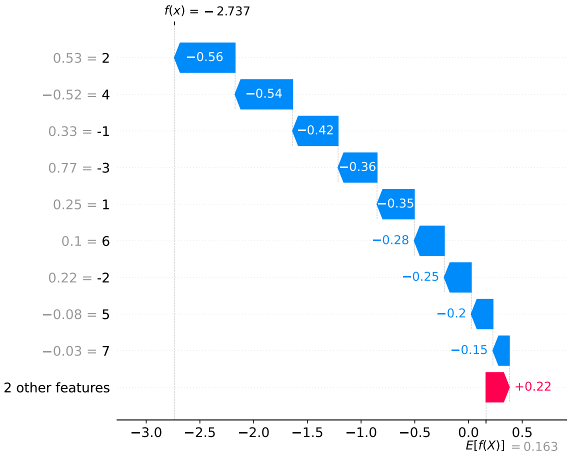


(u)

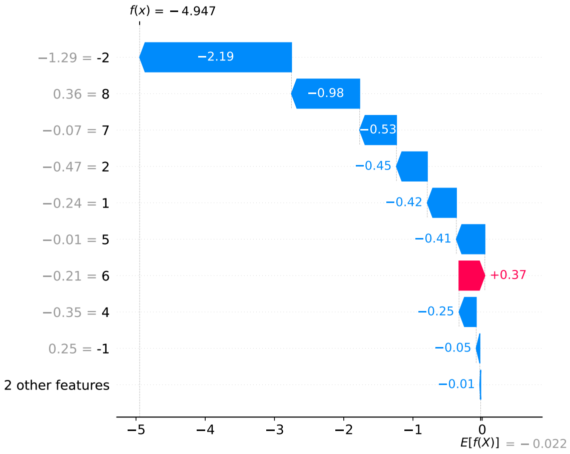


(v)

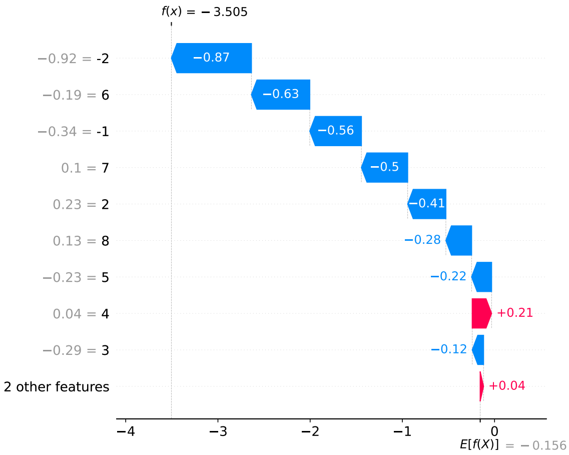


(w)

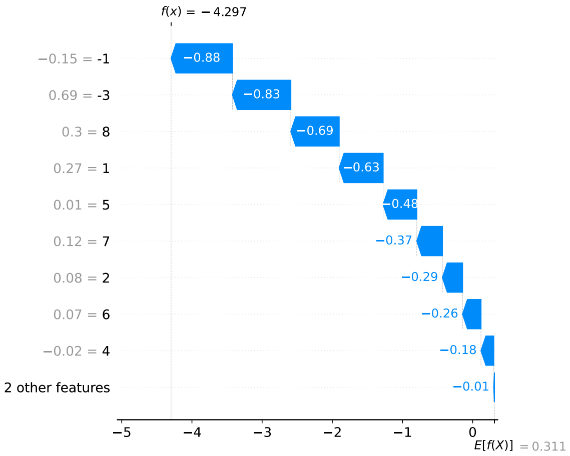


(x)

Fig. S3: (u–x: H3K79me, H3K9ac, H4K12ac, H4K5ac) SHAP waterfall plots for histone modifications — yeast gene *AAH1*.

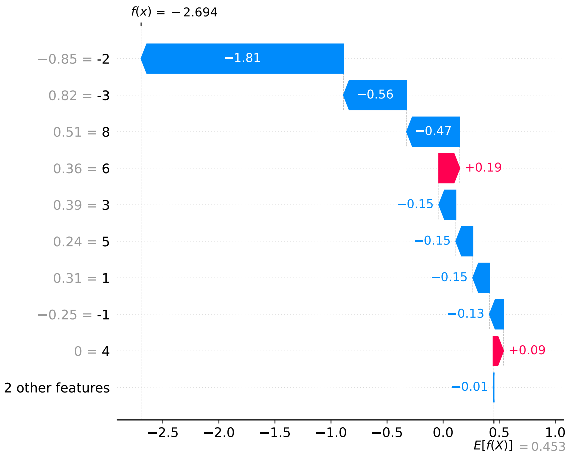


(y)

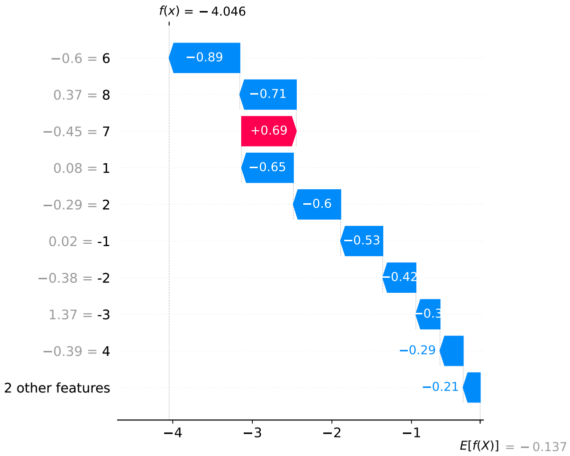


(z)

Fig. S3: (y–z: H4K8ac, Htz1) SHAP waterfall plots for histone modifications — yeast gene *AAH1*.

### 1.5 Stacked bar plots - Genebody PTMs

Stacked bar plots show average SHAP values across genebody positions. Blue bars indicate contributions toward higher expression, while orange bars indicate contributions toward lower expression. The distribution across positions highlights how activating and repressive marks shape expression patterns, and the SHAP magnitude reflects the importance of each PTM at specific positions.

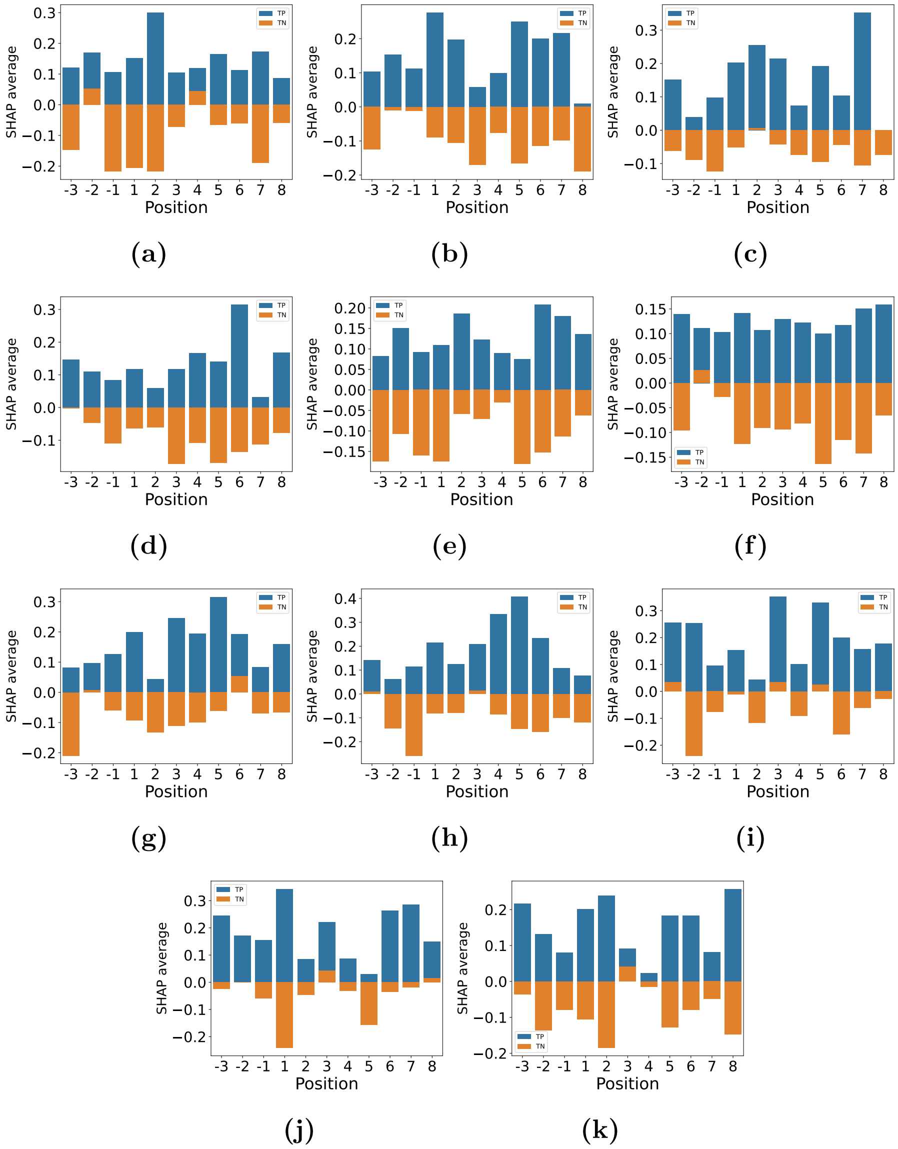


Fig. S4.Representative genebody modifications. Each panel (a–k: H3K4me, H4K20me, H3K79me3, H3K36me3, H4R3me2s, H4R3me, H3K36me, H3K36me2, H4K16ac, H3S10ph, H2AS129ph) shows one histone PTM.

### 1.6 Stacked bar plots – Promoter

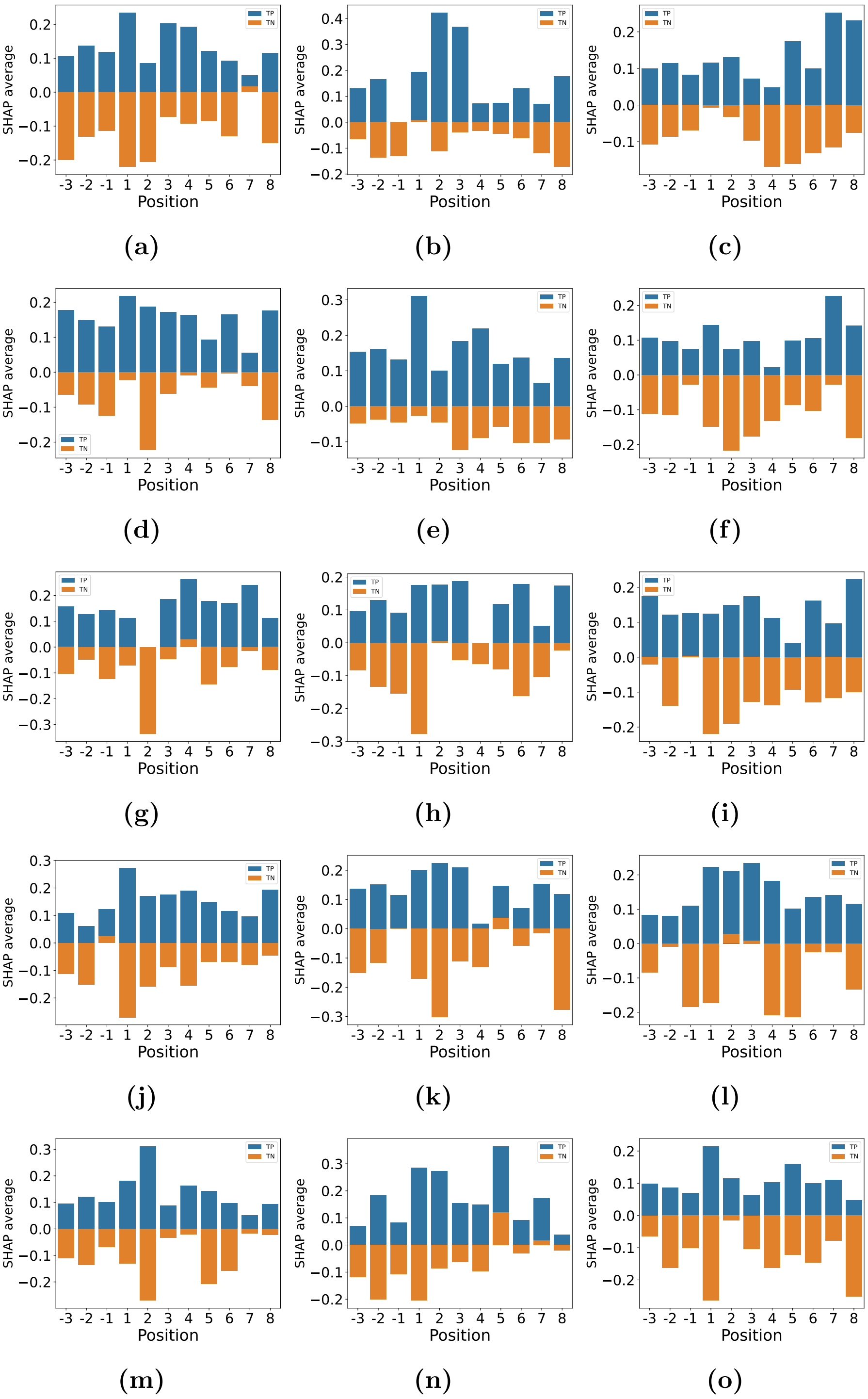


Fig. S5 Representative promoter modifications. Each panel (a–0: H2AK5ac,H3K14ac,H3K18ac,H3K23ac,H3K27ac,H3K4ac,H3K4me2,H3K4me3,H3K56ac,H3K79me,H3K9ac,H4[illegible] shows one histone PTM.

### 1.7 Beeswarm plots - Genebody

Beeswarm plots were used to visualize the contribution of gene-body histone modifications to transcriptional regulation. Each point in the plot represents a gene, with the SHAP value indicating the magnitude and direction of a modification's influence on the predicted transcriptional state. The spread of points reflects the variability in contribution across genes, while the color scale denotes the underlying modification intensity. These plots highlight which gene-body modifications consistently promote or repress transcription and reveal heterogeneity in their effects, emphasizing the positional importance of histone marks within the gene body

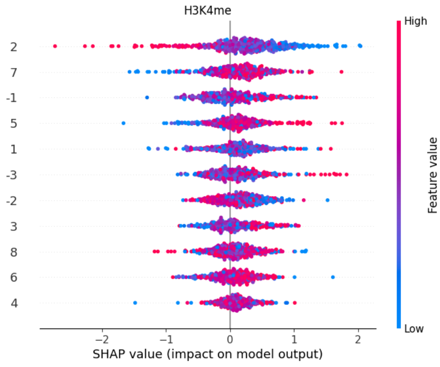


**(a)**

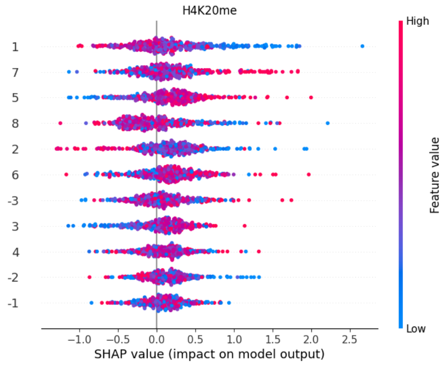


**(b)**

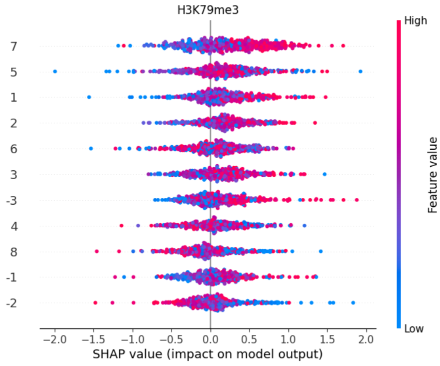


**(c)**

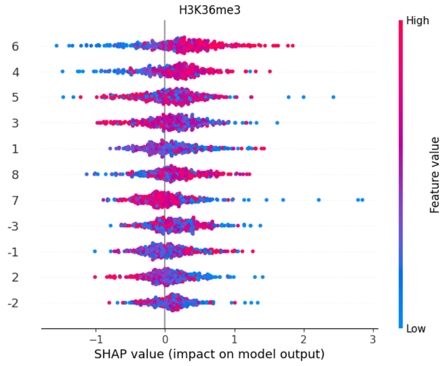


**(d)**

Fig S6 Beeswarm plots for representative genebody modifications (a–d : H3K4me,H4K20me,H3K79me3,H3K36me3)

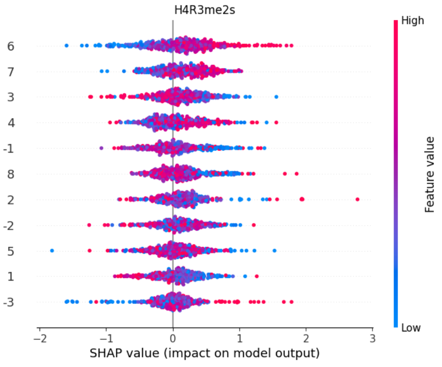


(e)

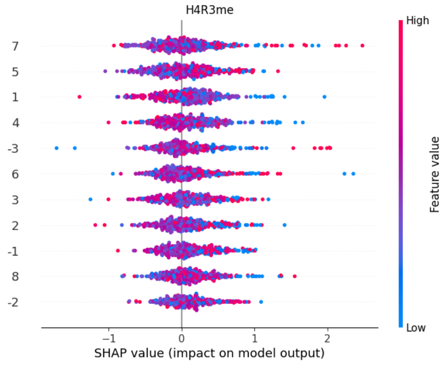


(f)

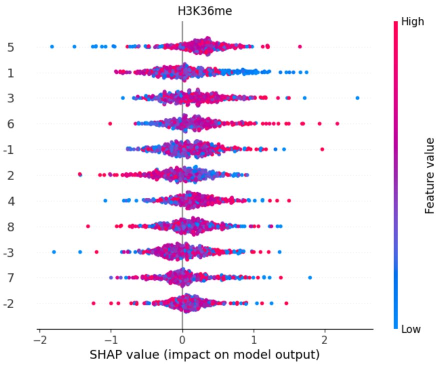


(g)

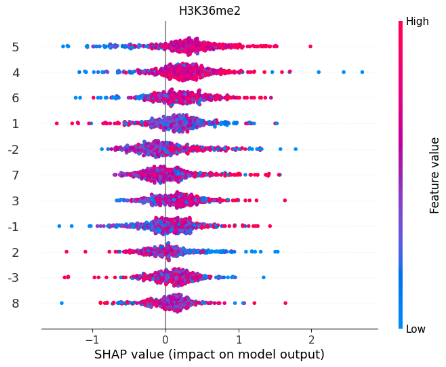


(h)

Fig S6 Beeswarm plots for representative genebody modifications (e–h : H4R3me2s,H4R3me,H3K36me,H3K36me2)

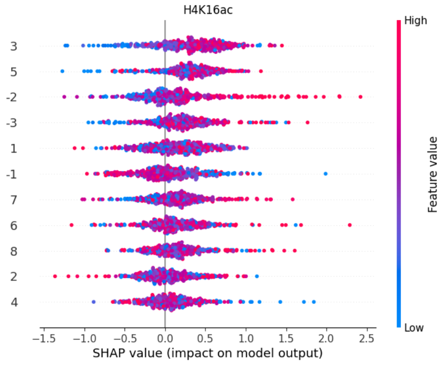


(i)

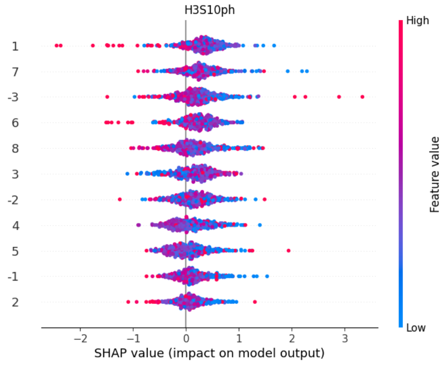


(j)

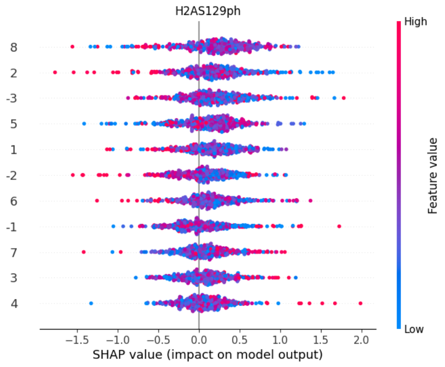


(k)

Fig S6 Beeswarm plots for representative genebody modifications (i–k : H4K16ac,H3S10ph,H2AS129ph1)

### 1.8 Beeswarm plots - Promoter

Beeswarm plots were used to illustrate the contribution of promoter-associated histone modifications to transcriptional regulation. Each point represents an individual gene, with SHAP values capturing the direction and strength of a modification's effect on the predicted transcriptional state. The distribution of points reflects gene-to-gene variability, while the color gradient indicates modification intensity at promoter positions. These plots reveal key promoter histone marks that strongly influence transcription initiation and highlight their position-specific regulatory roles.

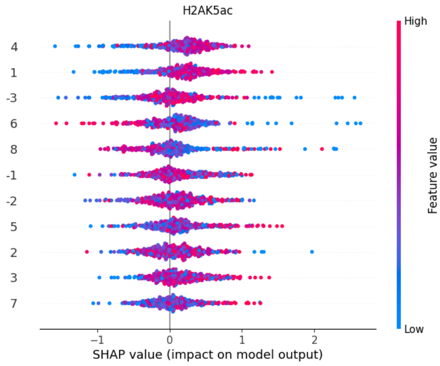


**(a)**

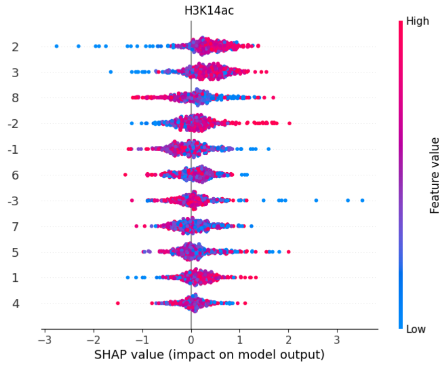


**(b)**

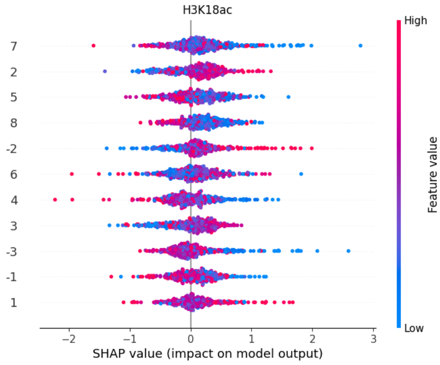


**(c)**

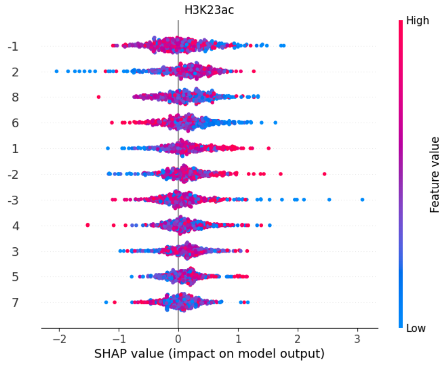


**(d)**

Fig S7 Beeswarm plots for representative promoter modifications (a–d : H2AK5ac, H3K14ac, H3K18ac, H3K23ac)

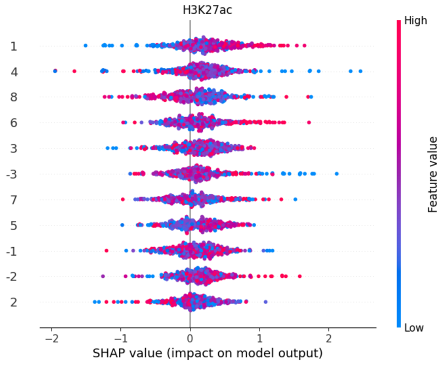


**(e)**

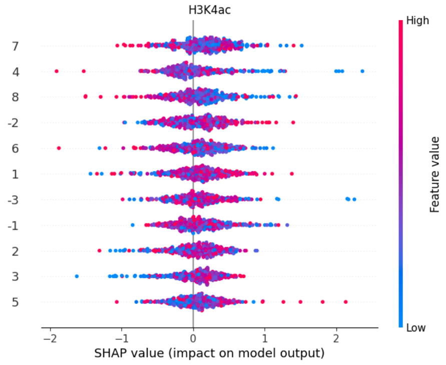


**(f)**

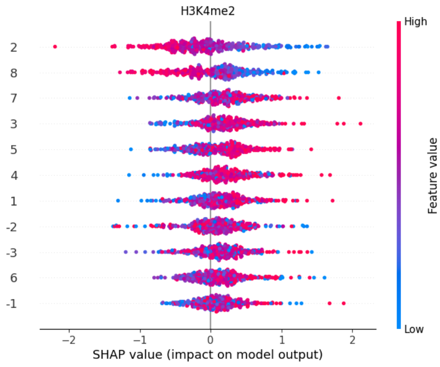


**(g)**

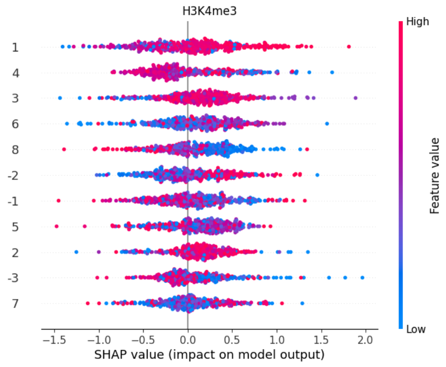


**(h)**

Fig S7 Beeswarm plots for representative promoter modifications (e–h : H3K27ac, H3K4ac, H3K4me2, H3K4me3)

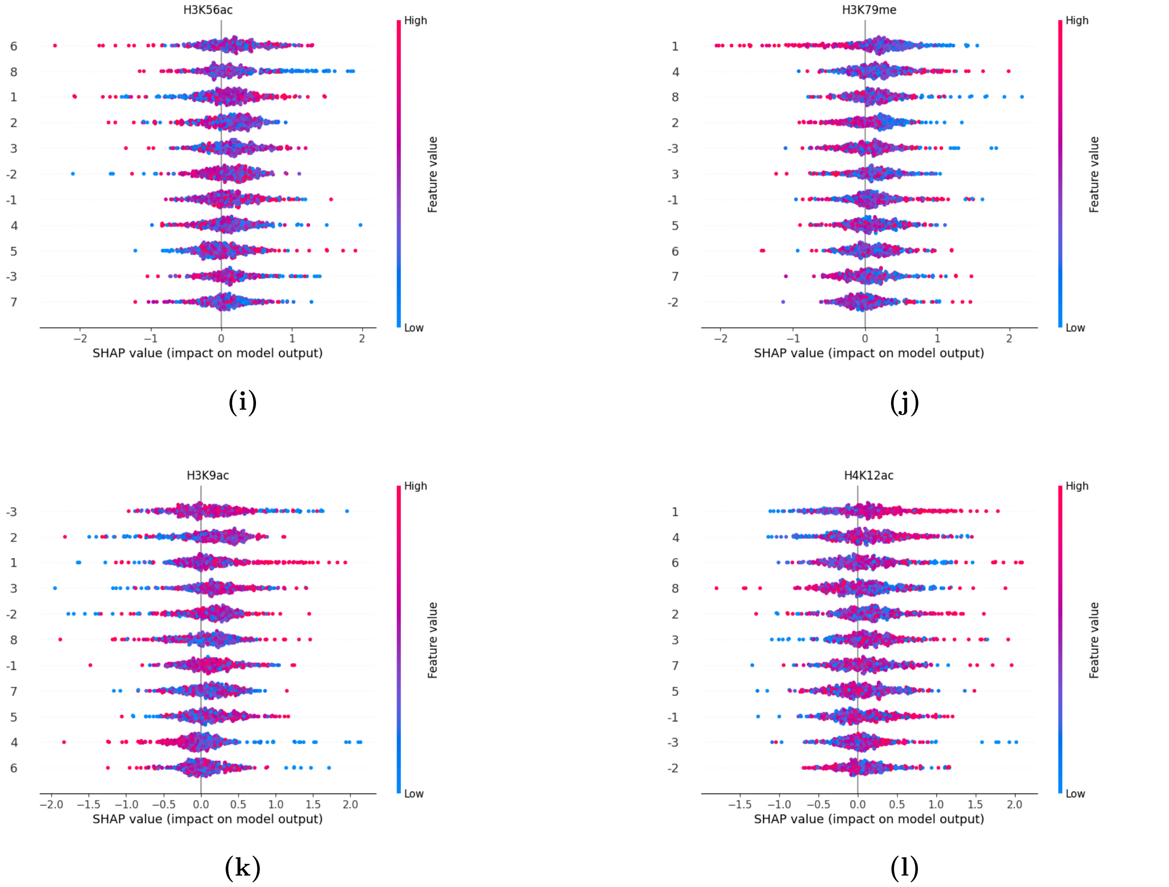


Fig S7 Beeswarm plots for representative promoter modifications (i–l : H3K56ac, H3K79me, H3K9ac, H4K12ac)

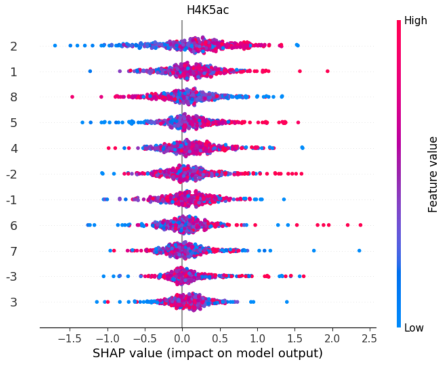


**(m)**

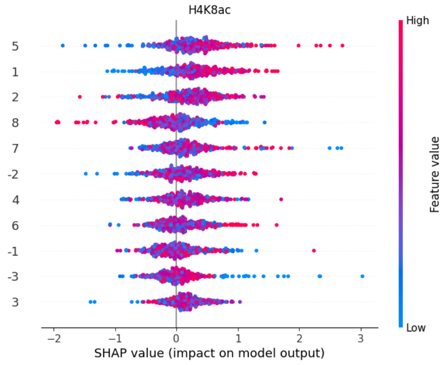


**(n)**

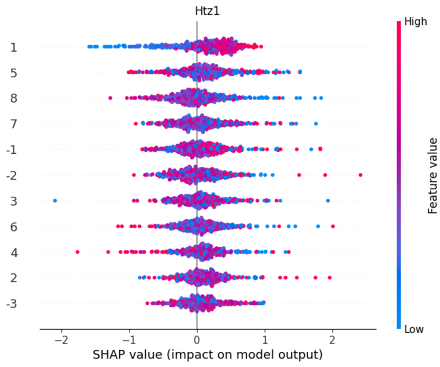


**(o)**

Fig S7 Beeswarm plots for representative promoter modifications (m–o : H4K5ac, H4K8ac, Htz1)
